# Taccalonolide AJ uncouples tubulin GTP hydrolysis from microtubule depolymerization

**DOI:** 10.64898/2026.09.30.755716

**Authors:** Ezekiel C. Thomas, Nicholas Vangos, Lauren A. McCormick, T. Lynne Blasius, Kristen J. Verhey, April L. Risinger, Michael A. Cianfrocco

## Abstract

Microtubules are dynamic polymers of tubulin subunits that are vital for a variety of eukaryotic cell functions, including intracellular trafficking and cell division. Microtubule-targeting drugs are among the oldest and most widely used chemotherapeutics. Despite their extensive use, the development and optimization of these drugs is hindered by our limited understanding of the molecular impacts on microtubule structure and function. Further, the widespread use of taxanes has led to taxane-resistant tumors, necessitating the development of potent microtubule targeting agents with distinct mechanisms of action. Recent high-resolution structures have begun to explain how paclitaxel (Taxol) stabilizes microtubules: it reverses the hydrolysis-driven lateral strain at the microtubule seam and induces an expanded, GTP-like conformational state that promotes polymerization. In this work, we demonstrate that the microtubule-targeting natural product taccalonolide AJ (tacca AJ) stabilizes microtubules in a manner completely distinct from paclitaxel despite binding to the same taxane site of β-tubulin. Using cryo-EM, we show that tacca AJ stabilizes a compacted, post-GTP hydrolysis state, which is usually prone to depolymerization. *In vitro* assays show that tacca AJ-stabilized microtubules are more rigid than paclitaxel-stabilized microtubules, and reconstituted dynamics with tacca AJ show increased rescue near the plus end. Finally, we use fluorescent biosensors for the compacted tubulin conformation to confirm that tacca AJ-stabilized microtubules in cells are compacted yet stable against depolymerization. We propose that the novel mechanism by which tacca AJ stabilizes microtubules underlies their efficacy in taxane-resistant tumor models and may serve as an entry point to development of next-generation chemotherapeutics.

## Introduction

Microtubules are essential eukaryotic cytoskeletal elements that support and drive fundamental biological processes including cell division, fast axonal transport, cell migration, and structural support within cilia and the mitotic spindle. Microtubules are composed of GTP-binding αβ-tubulin heterodimers. In the GTP-bound state, tubulin self-associates into elongated, hollow tubes^1^. After polymerization and within the microtubule lattice, β-tubulin hydrolyzes GTP to GDP. As long as the GTP-bound β-tubulin is present at the end of the microtubule - a ‘GTP-cap’ - the microtubule will continue to grow. However, if the plus end of the microtubule lattice contains GDP-β-tubulin, the microtubule will undergo a ‘catastrophe’ and depolymerize, shrinking in length. The process of microtubule growth and shrinkage is a hallmark of microtubule biology and is termed ‘dynamic instability^2^.’

Assembled microtubule lattices undergo structural changes that accompany nucleotide hydrolysis. When the microtubule consists of GTP-β-tubulin, the tubulin subunits within the lattice adopt an expanded conformation^3–5^. Upon GTP hydrolysis, the microtubule shifts to a compacted state in which its length decreases by ∼3%^4,5^ and switch-like regions within tubulin alter, producing longitudinal changes within the microtubule^6^. Recently, we developed a novel approach to determine cryo-EM reconstructions of microtubules^6–8^. Our data show that the expanded conformation exhibits favorable contacts across the entire microtubule lattice, including the microtubule seam^6^, the asymmetric point of contact within a 13-protofilament microtubule^9,10^, thereby conferring stability to the GTP lattice. Notably, the post-GTP hydrolysis conformation causes significant structural changes at the microtubule seam, where weakened contacts contribute to a loss of stability for the microtubule lattice^6^. Thus, the expanded pre-GTP hydrolysis conformation enables polymerization, whereas the compacted post-GTP hydrolysis conformation leads to depolymerization.

Given the importance of microtubules for eukaryotic cellular physiology, they are the targets of natural products produced by plants and microorganisms as toxins^11^. Microtubule-targeting agents can disrupt cellular processes by either driving microtubule depolymerization (*e.g.*, vinca alkaloids) or by stabilizing them in their polymerized state (*e.g.*, taxanes), both of which abolish dynamic instability. The diverse effects of microtubule-targeting natural products on inhibiting proliferation and oncogenic signaling make them one of the most widely used and effective classes of chemotherapeutic agents for a wide range of solid tumors and hematological malignancies^12^.

The taxane class of microtubule stabilizers, including paclitaxel, is widely used in the frontline treatment of many aggressive solid tumors, including as backbone chemotherapy in combination with new targeted agents and immunotherapy^12^. Paclitaxel binds to β-tubulin within the microtubule lumen distal to the nucleotide-binding site,^13^ but there are conflicting models on the mechanism by which this binding stabilizes microtubules^4,14^. Using our newly developed approach for determining high-resolution microtubule structures by cryo-EM, we discovered that paclitaxel exploits allosteric switches to convert the compacted, GDP-tubulin lattice into a GTP-like state^6^. By inducing an expanded lattice, paclitaxel allows the microtubule seam to convert from the unstable staggered conformation present in the compacted GDP lattice into an expanded form through favorable contacts, thus stabilizing the microtubule^6^.

Despite their well-documented efficacy in the treatment of a wide array of cancers, all clinically approved microtubule-targeted drugs are susceptible to the development of drug resistance and dose-limiting toxicities that limit their prolonged use. Thus, the identification and characterization of novel compounds that employ this well-validated anticancer mechanism through a distinct mechanism of action is critical for the development of next-generation chemotherapeutics. The taccalonolides (taccas), isolated from *Tacca chantrieri* plant extracts, were identified in the 2000s as a structurally distinct class of microtubule stabilizers that retain efficacy in taxane-resistant cancer models^15,16^. Subsequent semisynthetic work led to the C22,23 epoxidation of the natural taccas to generate the potent taccas AJ and AF that irreversibly bind and stabilize microtubules through a mechanism of action distant from the taxanes^17,18^. Structural and mutational analyses demonstrate that a highly specific covalent interaction occurs between the C22,23 epoxide of the taccas and the Asp224 residue of β-tubulin (Asp226 in publications where the α/β-tubulin alignment numbering convention is used^13,19^) to impart this profound and irreversible microtubule stabilization and provide efficacy against taxane-resistant cancer models both in vitro and in vivo^18,20–25^.

However, despite a co-crystal structure, hydrogen-deuterium exchange mass spectrometry (HDX-MS), mutational analysis, and modeling studies, there are conflicting models regarding the exact mechanism by which tacca AJ binds to and stabilizes microtubules. X-ray crystallography studies of tacca AJ bound to short tubulin oligomers stabilized in complex with the microtubule binding proteins stathmin and tubulin tyrosine ligase confirmed that the C22,23 epoxide of tacca AJ engages in a covalent interaction with β-tubulin Asp224, but found that this was associated with M-loop stabilization, in contrast to HDX-MS studies where this stabilization was only seen with taxanes, not the taccas^18,24^. Additionally, our mutational analysis and tacca structure activity relationship demonstrated that the interaction between Lys19 in β-tubulin and the C25 hydroxyl on the taccas is critical for the covalent binding site, despite the crystal structure suggesting this residue as a minor interaction^22,24,26^. Finally, in contrast to the GTP-bound form observed in the co-crystal structure, subsequent X-ray fiber diffraction showed that the microtubule longitudinal spacing was in a post-GTP hydrolysis, GDP-like conformation^27^. Thus, controversy remains over whether tacca AJ binds to pre-or post-GTP hydrolysis microtubule lattices, the critical contacts that mediate binding, and how this binding stabilizes microtubules to provide a spectrum of biological activity that differs from the taxanes.

Herein, we determined the mechanism by which tacca AJ stabilizes microtubules using a combination of cryo-EM with biochemical and cell-based assays. Our results unequivocally demonstrate that tacca AJ binds to a compacted, GDP-bound conformation of tubulin within intact microtubules and alters lateral interactions between tubulin subunits through a mechanism that corroborates our prior HDX-MS and mutational studies. We find that tacca AJ-stabilized microtubules are more rigid and are resistant to spermine treatment, supporting the strengthened lateral interactions observed in our structural analysis. Further, tacca AJ-stabilized microtubules display fewer catastrophes, slower depolymerization rates, and more frequent rescue events than the taxanes. Taken together, our data show that tacca AJ uncouples GTP hydrolysis from microtubule stability, preventing depolymerization of GDP-bound microtubules and providing a mechanistic understanding for a novel mechanism of pharmacologically induced microtubule stabilization.

## Results

To understand the molecular basis for tacca AJ-mediated stabilization of intact microtubules, we used single-particle cryo-EM analysis of microtubules covalently bound to tacca AJ. We leveraged our new analysis approach^6^ to break the microtubule pseudosymmetry, ultimately determining a 2.8 Å structure of a single protofilament from within a 13-protofilament microtubule (**Figure 1A & 1B**). The reconstruction showed strong density for tacca AJ bound to the taxane site of β-tubulin (**Figure 1B**). Furthermore, the cryo-EM density clearly showed that tacca AJ was bound covalently to β-tubulin, as indicated by the strong density connection (**Figure 1C**).

**Figure 1.**
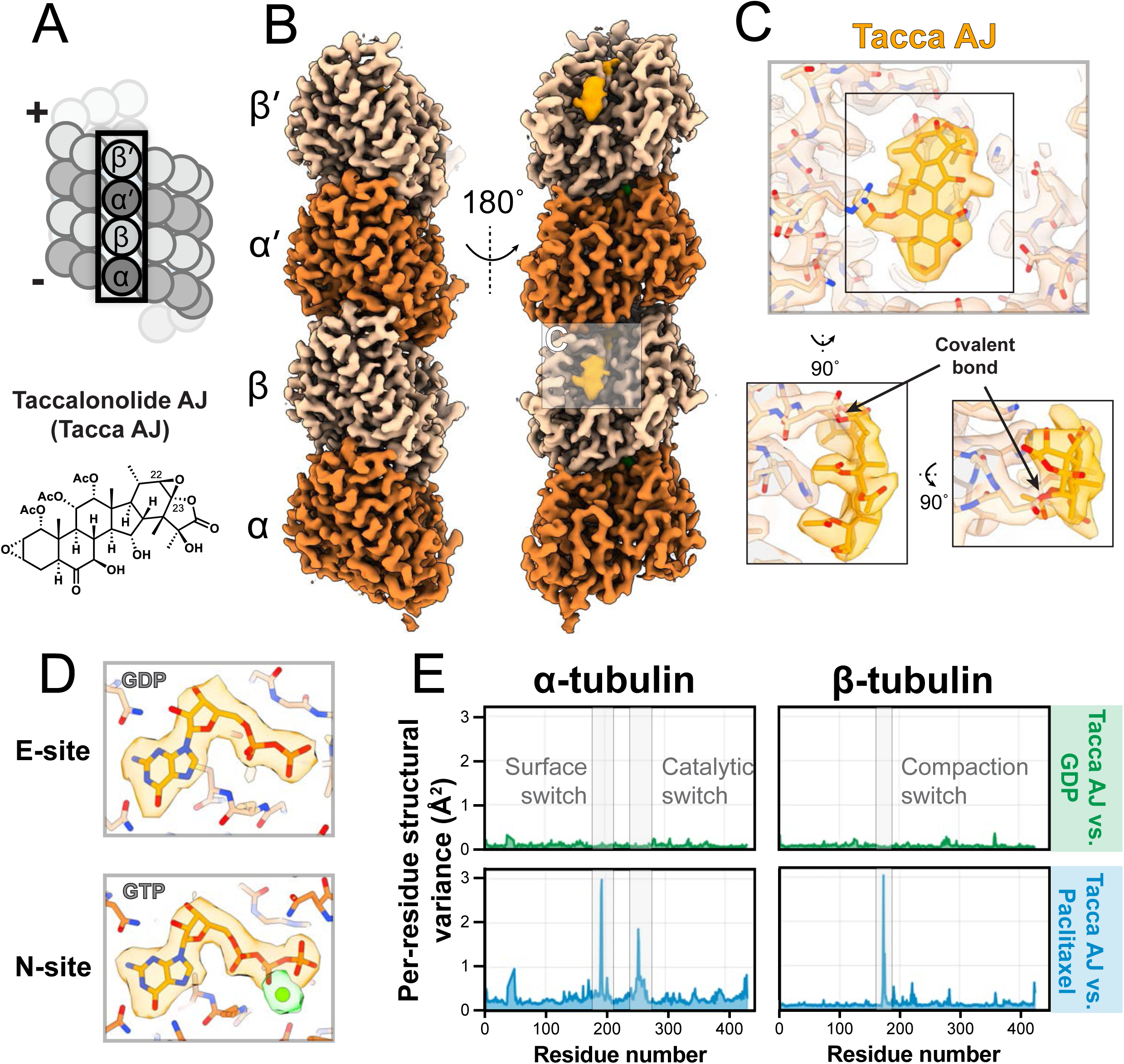
Tacca AJ-bound microtubules are GDP-bound and in a compacted conformation. (A) Schematic highlighting a single protofilament of a microtubule that was extracted and used for single particle cryo-EM reconstruction. (Bottom) Structure of taccalonolide AJ. (B) Cryo-EM reconstruction of a single protofilament. (C) Close-up view of the β-tubulin taxane site showing the atomic model within the cryo-EM density. (D) View of exchangeable (E)- and non-exchangeable (N)-site nucleotides with atomic model docked into cryo-EM density. Green density indicates magnesium density in N-site. (E) Per-residue structural variance calculated between tacca AJ-GDP vs. GDP and tacca AJ-GDP vs. paclitaxel-GDP.

Given conflicting prior data on the tubulin nucleotide state and lattice conformation of tacca AJ-tubulin, we next inspected the exchangeable nucleotide within β-tubulin (“E-site”). Our cryo-EM reconstruction clearly shows density corresponding to GDP, not GTP, in the E-site of β-tubulin (**Figure 1D**). This is further corroborated by comparing the non-exchangeable nucleotide site (“N-site”), which is constitutively bound to GTP, with the E-site, which is unambiguously GDP (**Figure 1D**).

Next, we compared the overall tubulin conformation in the tacca AJ-GDP reconstruction with our previous structures^6^ of GDP and paclitaxel-GDP microtubules using our per-residue structural variance measure, which provides a reference-free comparison of tubulin structures (**Figure 1E**). This comparison showed that the C-α positions of tacca AJ-GDP tubulin were nearly identical to those of GDP tubulin (**Figure 1E**). Further, comparison of tacca AJ-GDP tubulin with paclitaxel-GDP tubulin showed specific regions in paclitaxel-GDP that are in different conformations than tacca AJ-GDP (**Figure 1E**). These regions were previously identified as structural switches that change during GTP hydrolysis and are exploited by paclitaxel to expand the lattice conformation^6^. Thus, tacca AJ binds GDP-bound tubulin, and the microtubule lattice is in a post-GTP-hydrolysis, compacted conformation.

Comparison of the binding modes of tacca AJ versus paclitaxel shows that these microtubule-binding compounds use completely distinct modes of interaction. Both tacca AJ and paclitaxel bind to the taxane site of β-tubulin, which is a pocket found in β-but not α-tubulin (**Figure 2A & 2B**). Tacca AJ sits directly above β-tubulin helix 7, covalently bound to βAsp224, whereas paclitaxel interdigitates several moieties across the taxane pocket, ultimately sandwiching βHis227 (**Figure 2B**). Analysis of tacca AJ’s non-covalent interactions with β-tubulin shows that the only prominent interaction occurs on βLys19 (**Figure 2C**), consistent with previous mutagenesis studies showing that βLys19 was critical for tacca AJ binding to tubulin^22^. Superposition of tacca AJ and paclitaxel shows their unique binding modes, where tacca AJ remains closer to the charged surface of β-tubulin and paclitaxel binds deeper within the hydrophobic pocket (**Figure 2D**). Closer inspection of the interaction modes shows that the M-loop involved in lateral tubulin interactions is in the same conformation across GDP, tacca AJ-GDP, and paclitaxel-GDP tubulins (**Figure 2E & 2F**), in contrast to the previous crystal structure of tacca AJ bound to stathmin-tubulin, which proposed that tacca AJ contacted the M-loop^24^. Finally, further support regarding the distinct modes of tubulin interaction for tacca AJ and paclitaxel is shown via the nature of their interaction with βAsp224 and βHis227. Tacca AJ-bound β-tubulin shows that βAsp224 and βHis227 are in nearly the same conformation as GDP tubulin (**Figure 2E**), whereas paclitaxel utilizes a multivalent interaction with βHis227 to rotate its conformation, ultimately allowing a structured water molecule to bind between βHis227 and βAsp224, pushing βAsp224 away from paclitaxel towards the nucleotide site of β-tubulin (**Figure 2F**).

**Figure 2.**
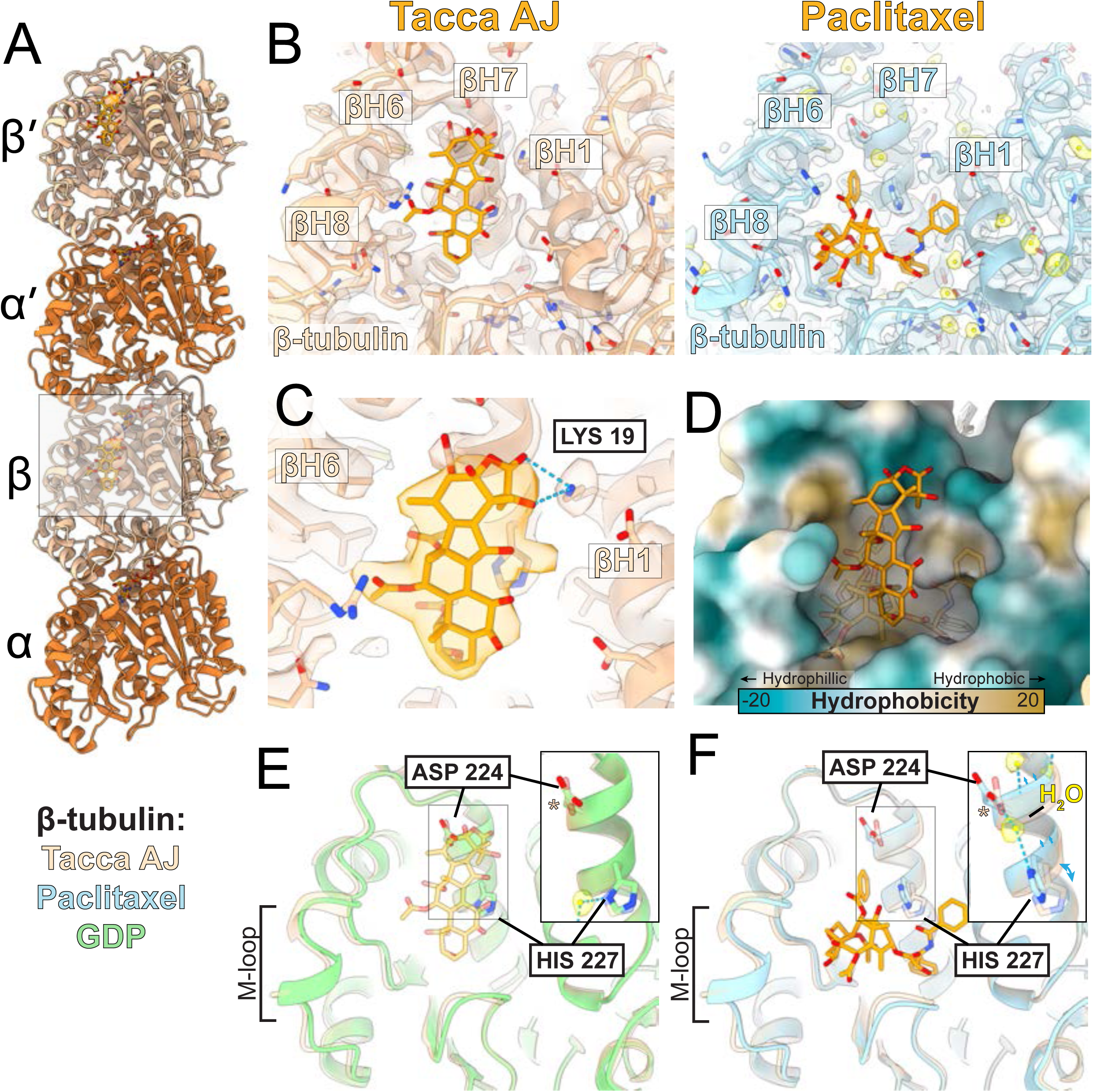
Tacca AJ binds to β-tubulin in a manner distinct from paclitaxel. (A) Overview of single protofilament atomic model of tacca AJ-GDP. (B) Comparison of tacca AJ-GDP model and density with paclitaxel-GDP^6^. Ligand density for tacca AJ and paclitaxel is hidden. Semi-transparent density is shown for β-tubulin. Water molecules from paclitaxel-GDP structure are highlighted in yellow. (C) Non-covalent interactions between tacca AJ and β-tubulin are shown, indicating that only βLys19 is proposed to make hydrogen bonds with tacca AJ. (D) Superposition of tacca AJ (solid) and paclitaxel (semi-transparent) is shown on β-tubulin hydrophobicity surface. (E) Superposition of tacca AJ-GDP (semi-transparent) with GDP^6^ (solid) model. Inset: close-up of βH7 with a structured water molecule from GDP shown in yellow. (F) Superposition of tacca AJ-GDP (semi-transparent) with paclitaxel-GDP^6^ (solid) model. Inset: closeup of βH7 with a structured water molecule from paclitaxel-GDP shown in yellow.

Zooming out from a single protofilament, we next investigated whether tacca AJ induced any lateral changes in the microtubules. We used our analysis approach^6^ to determine two protofilament structures from homotypic and heterotypic microtubule lattices (**Figure 3A & 3B**). Comparing the two protofilament tacca AJ-GDP structures with our previously determined GDP structures showed minimal lateral differences, consistent with our previous conclusion that tacca AJ-GDP exists in a compacted, GDP conformation (**Extended Data Figure 1**). However, we noticed one location on α-tubulin that showed a significant difference, prompting further investigation. Surprisingly, our homotypic lattice structure for tacca AJ-GDP showed that the H1-S2 loop, also known as the ‘α-tubulin K40 loop’ because it contains lysine 40 within α-tubulin, which can be acetylated^28^ or methylated^29^, becomes ordered within the homotypic tacca AJ-GDP lattice (**Figure 3C)**. This loop has never been observed in any other cryo-EM structures of undecorated microtubules. Notably, when this loop is acetylated, it makes the microtubule softer and more pliable, which is proposed to facilitate a long lifetime for stable, acetylated microtubules in cells^30^. Interestingly, we only see the ordering of the αK40 loop in the homotypic lattice, but not the heterotypic lattice (**Figure 3D**). Thus, our tacca AJ-GDP structure shows that the majority of the tubulin structure is unchanged between GDP and tacca AJ-GDP, and the largest structural difference is the ordering of the αK40 loop in tacca AJ-GDP in the homotypic lattice.

**Figure 3.**
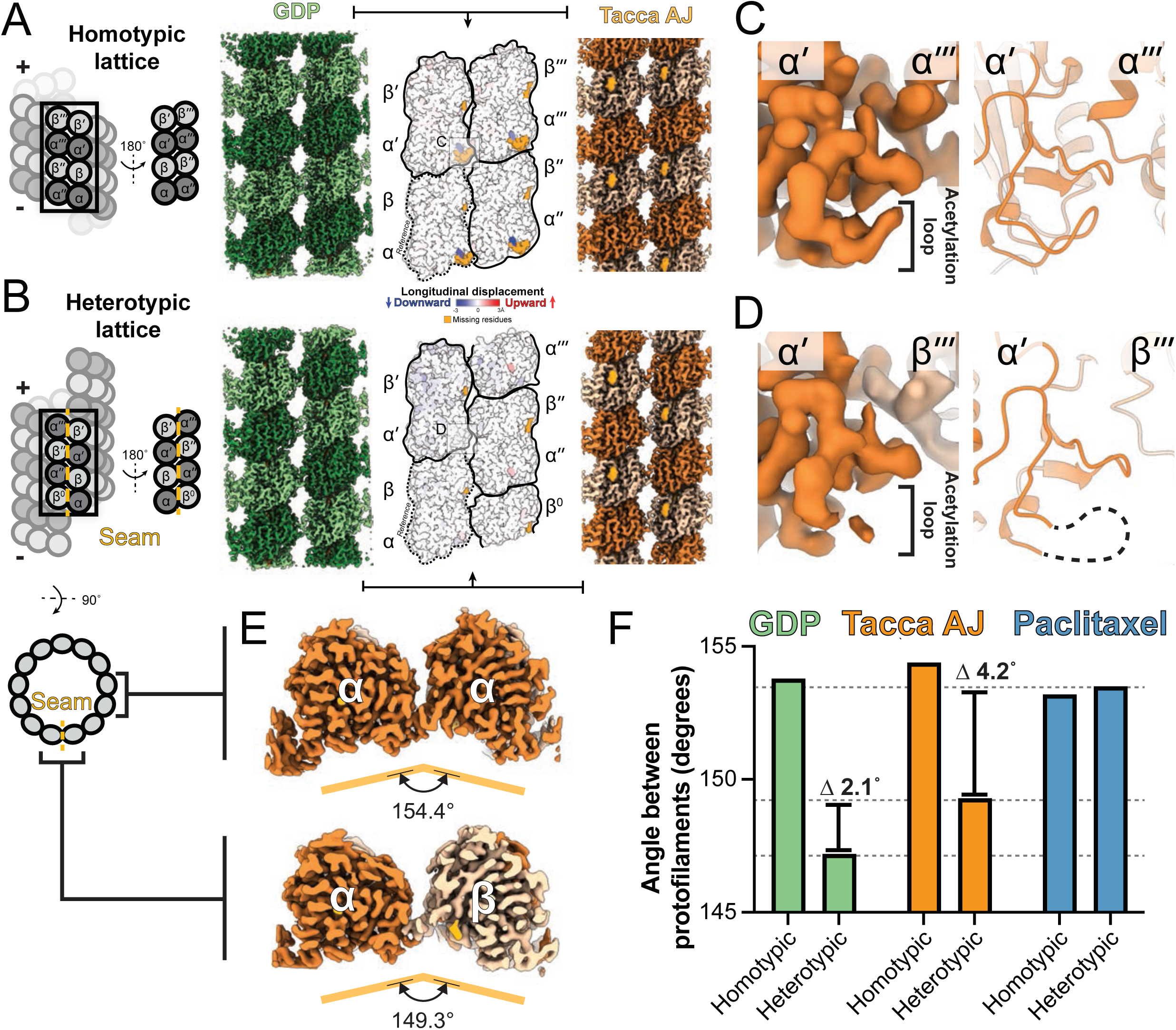
Tacca AJ induces lateral contact changes within the microtubule lattice. Homotypic (A) and heterotypic (B) lattices for two protofilament cryo-EM structures of tacca AJ-GDP are shown for comparison, highlighting longitudinal changes between GDP^6^ and tacca AJ-GDP homotypic lattices. Missing residues corresponding to the α-tubulin H1-S2 loop are colored orange. Close-up of α-tubulin density and atomic models from homotypic (C) and heterotypic (D) lattices. (E) Top-down view of two protofilament structures from homotypic (top) and heterotypic (bottom) lattices. (F) Comparison of interprotofilament angles for homotypic and heterotypic lattices for tacca AJ with GDP and paclitaxel-GDP^6^.

Next, we investigated the interprotofilament angle between the two protofilaments from the homotypic (2.8 Å) and heterotypic (3.1 Å) lattices (**Figure 3E**). Previously, we showed that GDP heterotypic lattices showed ‘staggered compaction’ and a narrower interprotofilament angle than the pre-GTP hydrolysis state^6^. This led us to model the GDP microtubule lattice as unstable due to structural changes at the microtubule seam. Comparison of the interprotofilament angle of tacca AJ-GDP with GDP and paclitaxel-GDP showed a slight decrease in the tacca AJ-GDP heterotypic lattice angle relative to GDP (**Figure 3E**). However, the overall angle of the heterotypic lattice in tacca AJ-GDP remained more similar to GDP than paclitaxel-GDP (**Figure 3F**, **Extended Data Figure 2**). These comparisons show small changes at the microtubule seam in tacca AJ-GDP, although, at this resolution, it remains challenging to infer whether this structural change is sufficient to induce a stronger microtubule seam than GDP.

Based on the altered lateral contacts observed in our cryo-EM structures, we predicted that tacca AJ microtubules would exhibit different physical properties *in vitro*. To measure the stiffness of the microtubules, we performed a gliding assay whereby actively transporting kinesin motors are adhered to the coverslip surface and microtubule velocity and movement behaviors can be observed. We thus compared the behavior of microtubules stabilized by the GTP-nucleotide analog GMPCPP, paclitaxel, or tacca AJ on coverslips coated with kinesin-1/KIF5C (**Figure 4A**). Microtubules analyzed in all three conditions had statistically similar lengths but exhibited differences in gliding velocity (**Extended Data Figure 3A & 3B**). Paclitaxel and GMPCPP microtubules showed faster gliding velocities (376 ± 83 and 473 ± 0 nm/s, respectively) than tacca AJ microtubules (329 ± 29 nm/s). Based on previous work^31^, these findings confirm that paclitaxel- and GMPCPP-stabilized microtubules are in an expanded state, given that kinesin-1 moves faster on expanded lattices, whereas the slower velocity of tacca AJ microtubules indicates they are compacted *in vitro*.

**Figure 4.**
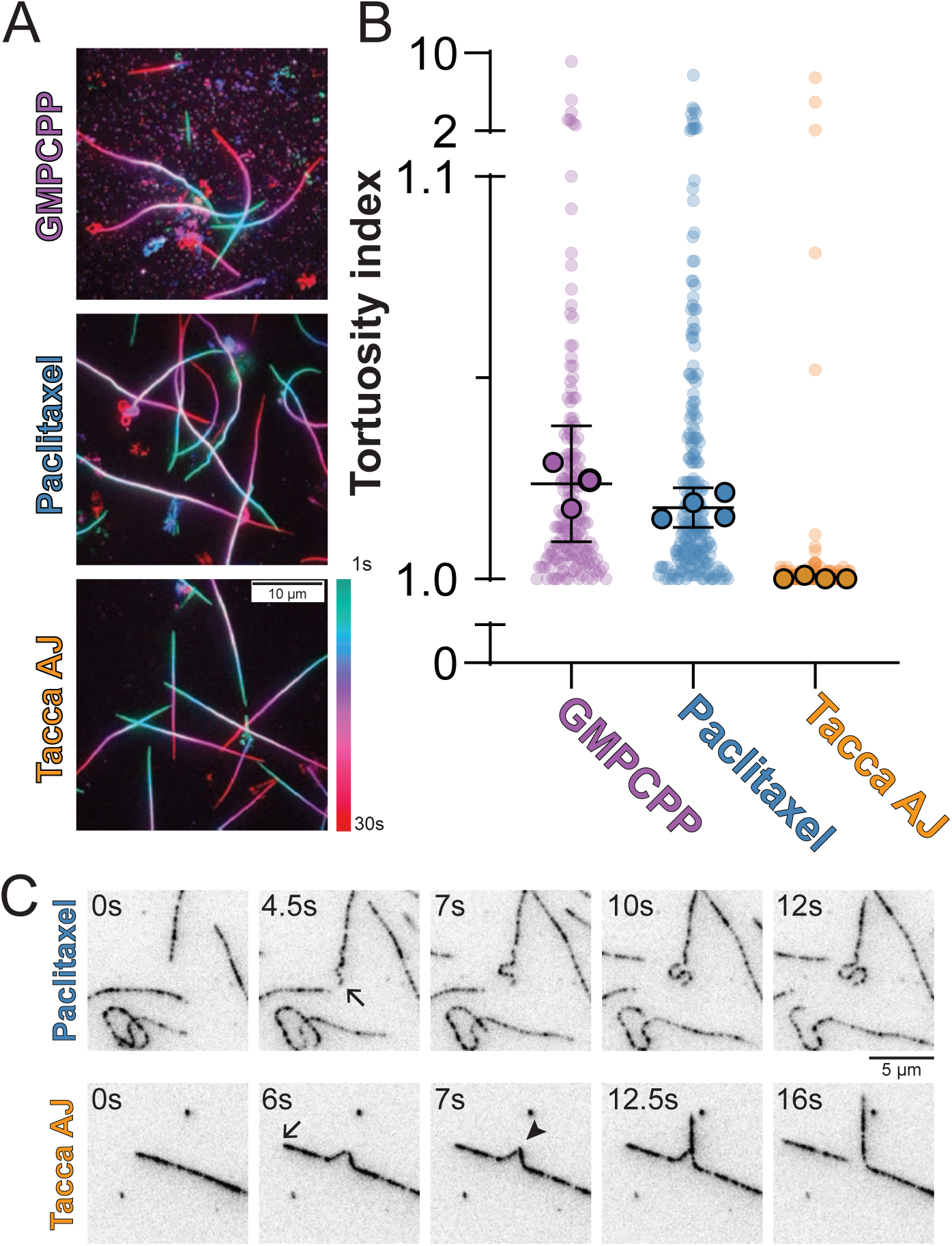
Tacca AJ-bound microtubules are rigid *in vitro*. (A) Example images of time-projected fluorescently labeled microtubules gliding along a kinesin-1-decorated coverslip. (B) Tortuosity index (arc-chord ratio) of microtubules from the first image of a gliding assay. Outlined points represent the median from each technical replicate. Error bars are the mean ± 95% confidence interval of technical replicates. GMPCPP n = 199 microtubules; paclitaxel n = 256 microtubules; taccalonolide AJ n = 206 microtubules. Adjusted p < 0.001 for taccalonolide AJ compared to GMPCPP or paclitaxel by one-way ANOVA with Tukey’s multiple comparisons test. (C) Example time points of paclitaxel- or taccalonolide AJ-stabilized microtubules (inverted greyscale) hitting unseen obstacles (arrows) in gliding assays. The paclitaxel microtubule bends but does not break. The taccalonolide AJ microtubule breaks at the point of greatest curvature (arrowhead). Scale bar = 5 µm.

Strikingly, whereas paclitaxel- and GMPCPP-stabilized microtubules displayed a propensity to bend while gliding and follow curving gliding paths, tacca AJ microtubules were rigid and followed straight gliding paths (**Figure 4A & 4B**). To quantitatively assess these different movement patterns, we measured microtubule tortuosity, the arc-chord ratio of the microtubules. A value of 1 indicates a straight line, and values >1 indicate a higher propensity to bend. Paclitaxel and GMPCPP microtubules had higher arc-chord ratios (1.018 ± 0.005 and 1.024 ± 0.014, respectively) compared to tacca AJ microtubules (1.000 ± 0.001). These results are consistent with paclitaxel increasing microtubule flexibility by altering lateral contacts, and our published structural data indicating paclitaxel- and GMPCPP-treated microtubules have similar lateral contacts. Further, these results indicate that tacca AJ microtubules are straighter and, likely, more rigid than paclitaxel- or GMPCPP-treated microtubules.

In our gliding assays, we noticed that tacca AJ microtubules were prone to breaking after being forced into a region of high curvature. This was evident when the leading end of the microtubule became anchored by an invisible obstacle (**Figure 4C, arrows**). For paclitaxel microtubules, the trailing microtubule adopted a highly curved morphology behind the anchor point without breaking. In contrast, the tacca AJ microtubules buckled behind the anchor point and would often break (**Figure 4C, arrowhead**), and then the newly formed microtubule end would continue to glide in a straight path. Furthermore, after breaking, the tacca AJ microtubules did not depolymerize, indicating the entire microtubule lattice has been stabilized by tacca AJ. This observation suggests that tacca AJ microtubules are more rigid than paclitaxel microtubules and resistant to depolymerization i*n vitro*.

Given the increased lateral contacts and rigidity of tacca AJ-stabilized microtubules, we hypothesized that tacca AJ microtubules would possess differences in end-stability compared to paclitaxel. High concentrations of the polyamine spermine have previously been shown to depolymerize paclitaxel-stabilized microtubules^32^. We predicted that the increased rigidity and altered lateral contacts of tacca AJ microtubules might resist spermine-induced microtubule depolymerization. We polymerized microtubules, stabilized them with paclitaxel or tacca AJ, and imaged them by Interference Reflection Microscopy (IRM) for 30 minutes after the addition of 30 mM spermine. Paclitaxel microtubules depolymerized slowly over this time period (24/24 microtubules showed depolymerization); however, tacca AJ-microtubules resisted spermine-induced depolymerization (only 1/50 microtubules displayed depolymerization) (**Extended Data Figure 3C**). Altogether, these results indicate that tacca AJ alters lateral contacts to stabilize microtubules in a unique mechanism that increases microtubule rigidity in a compact, post-GTP hydrolysis GDP lattice.

We next asked how the alteration of lateral contacts by tacca AJ would affect *in vitro* microtubule dynamics. Previous work using turbidity assays showed that unlike paclitaxel, tacca AJ does not lower the critical concentration for microtubule nucleation^18^, suggesting that tacca AJ stabilizes an existing microtubule lattice without affecting the interactions between soluble tubulin subunits.

To directly examine microtubule dynamics, we performed *in vitro* dynamics assays in which unlabeled microtubules were polymerized from stable GMPCPP seeds and dynamics were measured with IRM. In the control condition (vehicle-treated, ethanol), microtubules exhibited typical dynamics in which periods of growth ended in a catastrophe and depolymerization back to the stable GMPCPP seed (**Figure 5A**). In contrast, microtubules polymerized in the presence of 1 µM tacca AJ displayed frequent rescue events. Furthermore, we noticed that rescue events often occurred at the same site multiple times, then at a new rescue site closer to the dynamic plus end (**Figure 5A**).

**Figure 5.**
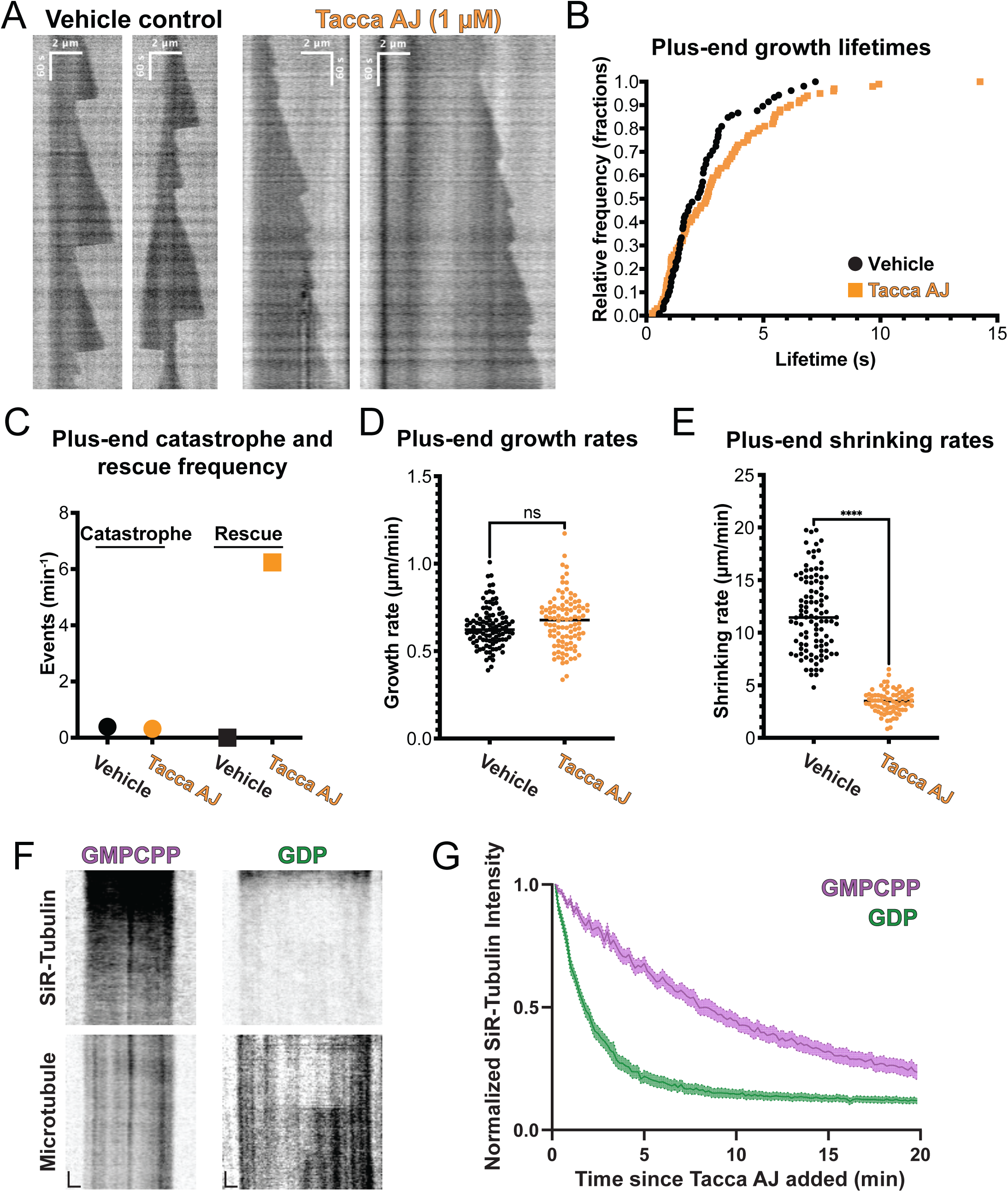
Tacca AJ incorporation into the GDP lattice promotes microtubule rescue. (A) Example kymographs depicting microtubule dynamics of unlabeled control or tacca AJ microtubules visualized by interference reflection microscopy. The more dynamic plus-end is oriented to the right of the images. (B) Cumulative relative frequency of plus-end growth lifetimes (vehicle n = 105 microtubules, tacca AJ n = 100 microtubules). (C) Catastrophe and rescue frequencies of microtubule plus-ends. (D) Growth rates of microtubule plus-ends. Bar = mean. p = 0.098 by Welch’s t-test. (E) Shrinking rates of microtubule plus-ends. Bar = mean. p < 0.001 by Welch’s t-test. (F) Kymograph showing SiR-Tubulin-bound GMPCPP (left) or GDP (right) microtubules after treatment with tacca AJ. Microtubules visualized by fluorescent tubulin. Scale bar height = 2 min, width = 1 µm. (G) Quantification of microtubule-bound SiR-Tubulin fluorescent intensity after displacement by tacca AJ on GMPCPP (n = 50 microtubules) or GDP microtubules (n = 49 microtubules). Shaded area = 95% confidence interval of the mean.

As a result, microtubules treated with tacca AJ showed increased plus-end growth lifetimes (**Figure 5B**) and higher rescue rates (vehicle = 0 rescues/min, tacca AJ = 6.24 rescues/min) (**Figure 5C**). Plus-end growth rates were not statistically different from control, consistent with a model in which tacca AJ preferentially acts away from the plus-end on a compacted, GDP-bound lattice (vehicle = 0.637 ± 0.02 µm/min, tacca AJ = 0.668 ± 0.03 µm/min) (**Figure 5D**). However, tacca AJ decreased plus-end shrinking rates (vehicle = 11.77 ± 0.71 µm/min, tacca AJ = 3.45 ± 0.24 µm/min), suggesting that the dynamic region near the plus end of the microtubule may represent a partially bound but insufficiently stabilized microtubule lattice (**Figure 5E**).

Given these effects on microtubule dynamics, we next wanted to confirm that tacca AJ incorporation into the lattice in our microtubule dynamics assay was within the body of the microtubule (GDP-bound tubulin) and not the plus end (GTP-bound tubulin). To test this, we compared the ability of tacca AJ to bind to expanded vs compacted lattices. As tacca AJ binding cannot be observed by fluorescence microscopy, we used the displacement of SiR-tubulin, a fluorescently labeled taxane derivative, as a proxy for tacca AJ binding, as these compounds compete for the taxane pocket. We generated expanded lattices by polymerization in the presence of GMPCPP and compacted lattices by polymerization in the presence of GTP, which is hydrolyzed to a compacted GDP-bound lattice. Upon addition of tacca AJ, the SiR-tubulin signal was lost ∼4-fold faster from compacted than expanded lattices (50% reduction in SiR-tubulin intensity after 2 min for GDP or 8 min for GMPCPP) (**Figure 5F & 5G**), which was not observed when treated with vehicle control (**Extended Data Figure 3)**.

These results agree with past observations and support a model in which tacca AJ preferentially acts on a compacted, GDP lattice rather than an expanded, GTP-like microtubule end. These stabilized GDP lattices then serve as regions that prevent depolymerization while also promoting new microtubule growth.

All our data thus far demonstrated that tacca AJ stabilized GDP lattices within *in vitro*microtubules. Next, we asked how tacca AJ affects microtubules in cells. First, we wanted to confirm that adding tacca AJ to cells influenced microtubule dynamics. Given that we did not have a fluorescent tacca AJ to confirm the timescale of tacca AJ binding, we instead monitored microtubule dynamics and the localization behavior of the plus-end tracking protein EB3, a marker for dynamic, growing microtubule plus-ends. We performed live cell imaging and measured the loss of EB3 comets in U2OS cells (**Extended Data Figure 4**). When treated with vehicle control, there was no change in the number of EB3 comets over the course of 20 minutes. When treated with 2.5 µM paclitaxel, the number of EB3 comets per cell was reduced by ∼75% within 1 minute, consistent with past results calculated using fluorescently labeled paclitaxel^33^ (**Extended Data Figure 4**). Notably, EB3 decorates the microtubule lattice at high expression levels, and paclitaxel treatment resulted in a rapid loss of lattice-bound EB3 within the 10-second imaging interval. With 2.5 µM tacca AJ, EB3 comets were lost more slowly than with paclitaxel, with a ∼50% reduction within 5 minutes, and most EB3 comets were lost by 10 minutes post-treatment (**Extended Data Figure 4**). Although tacca AJ reduced EB3 comets, microtubule lattice-bound EB3 remained. These results indicate that, at low micromolar concentrations, both paclitaxel and tacca AJ disrupt microtubule dynamics within minutes but affect the microtubule lattice differently.

Having established that tacca AJ disrupts microtubule dynamics, we also wanted to ensure that the microtubules in treated cells are stable and resist depolymerization. We transfected cells with GFP-α-tubulin to directly observe microtubules and performed a cold shock to test microtubule stability. We first treated cells for 30 minutes with 2.5 µM paclitaxel or tacca AJ, which did not significantly affect microtubules. Then, we cold-shocked the cells for 10 minutes at -20°C to promote microtubule depolymerization (**Figure 6A**). As expected, microtubules were completely depolymerized in vehicle-treated cells, showing a diffuse GFP signal (**Figure 6B**). In contrast, microtubules remained intact in paclitaxel- or tacca AJ-treated cells (**Figure 6B**). These results indicate that tacca AJ stabilizes microtubules in cells.

**Figure 6.**
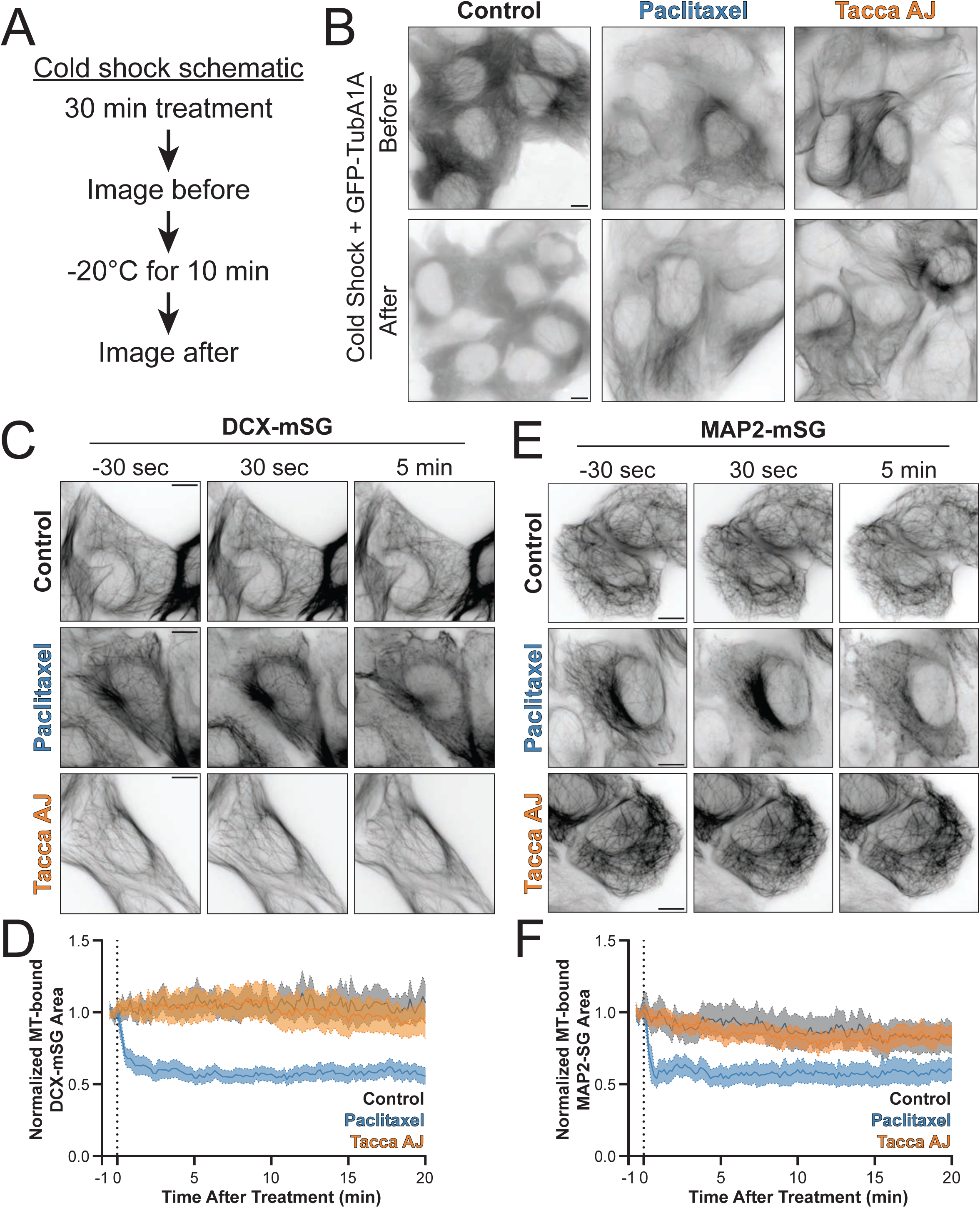
Tacca AJ stabilizes compacted microtubules in cells. (A) Schematic of GFP-α-tubulin cold shock assay. (B) Example images of GFP-α-tubulin in treated U2OS cells before and after cold shock. (C) Example time-point image of DCX-mStayGold intensity (inverted greyscale) in live U2OS cells after control, paclitaxel, or tacca AJ treatment. (D) Quantification of microtubule-bound DCX-mStayGold fluorescent intensity after treatment, normalized to intensity before treatment. Line = mean, shaded area = 95% confidence interval. Control n = 52 cells; paclitaxel = 76 cells; tacca AJ = 80 cells. (E) Example time-point images of MAP2-mStayGold intensity (inverted greyscale) in live U2OS cells after control, paclitaxel, or tacca AJ treatment. (F) Quantification of microtubule-bound MAP2-mStayGold fluorescent intensity after treatment, normalized to intensity before treatment. Line = mean, shaded area = 95% confidence interval. Control n = 36 cells; paclitaxel = 64 cells; taccalonolide AJ = 60 cells. All scale bars = 10 µm.

To determine whether tacca AJ maintains a compacted, GDP-like microtubule lattice in cells, we expressed DCX-mStayGold, which is sensitive to lattice conformation and preferentially binds to compacted GDP lattices^34–36^. Overexpression of DCX results in DCX microtubule localization throughout cells (**Figure 6C**). Adding 2.5 µM paclitaxel rapidly disrupts microtubule-bound DCX, leaving only fragmented microtubule-bound regions after several minutes (**Figure 6C & 6D**). In stark contrast, adding 2.5 µM tacca AJ has no effect on microtubule-bound DCX, comparable to the vehicle control (**Figure 6C & 6D**). To confirm that tacca AJ does not expand the microtubule lattice, we repeated these experiments using MAP2, which has a different microtubule-binding domain that also preferentially binds to a compacted lattice state^37^. Similar to the effects on DCX localization, paclitaxel quickly disrupted MAP2 microtubule binding, but tacca AJ maintained MAP2 localization comparable to vehicle control (**Figure 6E & 6F**). The rapid paclitaxel-driven loss of DCX and MAP2 from microtubules is consistent with its effects on microtubule dynamics; despite tacca AJ disrupting EB3 comets within a similar timeframe, DCX and MAP2 microtubule localization remained unchanged during this period. Nonetheless, to rule out the possibility that tacca AJ was slower to alter lattice conformation, we imaged cells for 12 hours after treatment. Over this longer period, both DCX and MAP2 microtubule binding was disrupted by paclitaxel but not by tacca AJ treatment (**Extended Data Figure 5**). Overall, these results suggest that, uniquely relative to paclitaxel, tacca AJ stabilizes the microtubule lattice in cells in a compacted conformation.

## Discussion

Here, we used cryo-EM to determine structures of tacca AJ covalently bound to β-tubulin and confirmed that the lattice is in a post-GTP hydrolysis state with a compacted conformation.

Despite binding to the same β-tubulin pocket, tacca AJ makes unique contacts relative to paclitaxel, and thus induces unique changes in lateral contacts. *In vitro* microtubule reconstitutions corroborated our structural findings and revealed that tacca AJ treatment yields rigid microtubules with stable rescue sites that stop depolymerization. Finally, in cell analyses confirm that tacca AJ quickly disrupts microtubule dynamics and stabilizes a compacted lattice conformation.

We propose a unique mechanism by which tacca AJ stabilizes microtubules in a compacted lattice conformation. Unlike paclitaxel, tacca AJ preferentially binds to a post-hydrolysis, GDP-bound microtubule lattice, which explains why tacca AJ does not lower the critical concentration required for microtubule nucleation; therefore, microtubule nucleation and polymerization are unaffected by tacca AJ. However, after tubulin is incorporated into the lattice and undergoes hydrolysis and compaction, tacca AJ binds and begins to stabilize the microtubule lattice, creating stable regions where rescue can repeatedly occur. As polymerization stochastically continues for uninterrupted periods, tacca AJ can bind more microtubule lattice and the tacca AJ-stabilized region “follows” the growing plus-end of the microtubule. Consequently, tacca AJ-treated microtubules display a short, dynamic microtubule region close to the plus-end that might represent a partially tacca AJ-bound but not yet fully stabilized region, and a fully tacca AJ-bound and stabilized bulk microtubule polymer.

Regarding the structural source of microtubule stability, tacca AJ is intriguingly in a conformation very similar to an unstable post-GTP hydrolysis state. Previous work suggests that paclitaxel induces microtubule stability by reversing molecular switches that occur during GTP hydrolysis^6^, at least in part by interacting with βHis227 in helix 7. Instead, tacca AJ covalently binds βAsp224 and sits above helix 7, which may help maintain a post-GTP hydrolysis, compacted conformation. The greatest structural difference relative to a GDP microtubule is the ordered nature of the α-tubulin H1-S2 loop, which has never been observed in an undecorated microtubule structure. Acetylation of αLys40 within this loop has been implicated in increased microtubule flexibility and reduced protofilament lateral interactions^30,38,39^; in contrast, tacca AJ results in more rigid microtubules, presumably due to increased lateral interactions. Importantly, we do not observe density for this loop in the two-protofilament heterotypic lattice, which has different lateral contacts. Furthermore, tacca AJ microtubules resist the chemically induced outward bending of protofilaments, preventing microtubule depolymerization. Overall, we propose that tacca AJ stabilizes microtubules by increasing lateral interactions and microtubule rigidity, which in turn prevents depolymerization.

An important future question remains: how subtle allosteric changes propagate through tubulin to yield dramatic differences in microtubule stabilization and function. Multiple classes of compounds target the β-tubulin taxane pocket but have opposite effects on lattice conformation, such as paclitaxel and tacca AJ. Differences in interactions with β-tubulin helix 7 may help explain some of these effects, but other covalently binding compounds modify β-tubulin helix 7 and have conflicting effects: zampanolide stabilizes a compacted conformation, whereas dactylolide stabilizes an expanded conformation^27^. High-resolution structures of these drugs bound to microtubules are needed to clarify these differences. Moreover, laulimalide and peloruside A are microtubule-stabilizing compounds that bind to a separate β-tubulin site that is closer to protofilament lateral contacts. Both of these compounds also stabilize a compacted conformation, but the specific mechanism remains uncertain. Our work demonstrates that increasing lateral interactions between protofilaments is a likely mechanism for stabilizing microtubules in a compacted conformation without reversing nucleotide-driven structural changes.

Our discoveries of the unique properties of tacca AJ-induced microtubule stabilization have important implications for potential clinical applications. All currently used microtubule-stabilizing drugs in the clinic induce an expanded conformation, likely because of how they stabilize microtubules; however, tacca AJ stabilizes a compacted conformation. The microtubule field is beginning to appreciate that these conformational differences can have wide-reaching cellular effects. Indeed, in our work reported here, tacca AJ uniquely affects the interactions with microtubule-associated proteins and kinesin-1 motors. Common mechanisms of taxane resistance involve changes in the expression of tubulin isotypes or microtubule-associated proteins. Tacca AJ remains effective in taxane-resistant cell lines, potentially because of these altered interactions.

## Acknowledgements

We thank Cianfrocco, DeSantis, McCormick, Sept, and Verhey labs for critical discussions of this work, especially Patrick DeLear for help with the Per Residue Structural Variance analysis. We acknowledge the following funding sources: S10OD030275 (N.V. & M.A.C), R01GM141119 (N.V. & M.A.C.), T32GM145470 (N.V.), R35GM131744 (K.J.V.), U54CA280805 (L.A.M.), and American Cancer Society PF-24-1320851-01-CCB (E.C.T.). This work was supported in part by the Greehey Distinguished Chair in Targeted Molecular Therapeutics (A.L.R.). This research utilized resources of the Structural Biology Core Facilities, part of the Institutional Research Cores at the University of Texas Health Science Center at San Antonio supported by the Office of the Vice President for Research and the Mays Cancer Center Drug Discovery and Structural Biology Shared Resource (NIH P30 CA054174) with the support of UT STARs awards 402-1288 and 402-1317.

## Materials & Methods

### Cryo-EM grid preparation and data collection

20 µM tubulin (Cytoskeleton) with three-fold molar excess of Taccalonolide AJ was prepared in 100 µL 80 mM PIPES pH 6.8, 1 mM EGTA, 1 mM MgCl_2_, 10% glycerol and 1 mM GTP on ice and warmed to 37 °C for an hour in a Cytation 5 plate reader (BioTek) where microtubule polymerization was confirmed by monitoring absorbance at 340 nm. Polymerized microtubules were centrifuged for 30 min at 20,000g at 37 °C and resuspended in 80mM PIPES pH 6.8, 1mM EGTA, 1mM MgCl_2_, without glycerol or GTP. For vitrification using a Vitrobot Mark IV (Thermo Fisher Scientific), 3 µl of sample was applied onto Au Quantifoil 3.5/1 200 mesh grids with a 2 nm ultrathin carbon coating (Electron Microscopy Sciences) at 37°C and 95% humidity. After a 30 second wait time, grids were subsequently blotted for 3.5 s with a blot force of 10 and plunged into the liquid ethane. Grids were transferred into liquid nitrogen and stored at cryogenic temperatures before clipping, screening, and data collection. Cryo-EM screening and data collection were conducted at the UT Health San Antonio Cryo-EM Facility on a Glacios TEM equipped with a Falcon 4 and Selectris energy filter. We recorded movies at a calibrated physical pixel size of 0.88838 Å/pixel with a total exposure of 52 e⁻/Å².

### Cryo-EM whole-microtubule data processing

Whole-microtubule processing was performed as previously described^6^ with minor modifications (**Extended Data Figure 6**). Imported movies were motion-corrected using RELION’s (v.5.0.0) own implementation of MotionCor2^40^. CTFFIND-4.1.14^41^ was used to estimate the CTF of each micrograph. Micrographs were filtered to better than 7 Å maximum resolution, yielding 9,242 micrographs. Particles were then picked using a trained neural network filament picker in crYOLO^42^. The resulting 1,209,568 particles were extracted at 4x binning with a 172 pixel (∼612 Å) box and subjected to a round of 2D classification, yielding 935,484 particles. 250,000 particles were randomly subsetted from this larger stack to make downstream analysis more computationally feasible.

Microtubule particles were subjected to a supervised 3D classification using 6 synthetic references to sort by protofilament number, as previously described^43^, yielding mostly 13-and 14-protofilament microtubules. This step resulted in 113,192 13-protofilament tacca AJ-GDP particles. These particles were then re-extracted unbinned with a 690 pixel (∼612 Å) box size and subjected to 3D refinement with applied helical symmetry (search range: -27.4° to -28.0°, 8.9 Å to 9.7 Å), yielding a 3.98 Å reconstruction. Particles were CTF-refined (per-micrograph defocus, per-micrograph astigmatism) and subjected to Bayesian polishing (default parameters). Subsequent 3D refinement with applied helical symmetry yielded a 3.54 Å reconstruction. Particles were then symmetry expanded by 26-fold to account for all possible subunit positions using the helical parameters from the final refinement jobs (**Table 1**), yielding 2,942,992 particles. Symmetry-expanded particles were subjected to signal subtraction with re-centering on a 400-pixel (∼355 Å) box using either a 1- or 2-protofilament mask, resulting in 2 signal-subtracted particle stacks. All subsequent 3D refinements were performed without applied symmetry.

### Cryo-EM 1-protofilament data processing

1- protofilament processing was performed as previously described^6^ (**Extended Data Figure 7, left**). 1-protofilament subtracted particles were backprojected to generate reference maps for subsequent refinements. Initial 3D refinement yielded a 3.12 Å map with a blurred α-tubulin and β-tubulin register. To sort the tubulin register, 2 high-resolution (6 Å) reference maps were generated using the molmap command in ChimeraX^44^ by fitting human tubulin (pdb: 5IJ0) into the initial mixed-register map in both possible lattice configurations. Refined particles were then subjected to a supervised 3D classification without alignment, using the previous 2 molmaps as unfiltered 6 Å references and a regularisation parameter T of 200. This step leverages high-resolution prior information, which is the reason for the high T value and unfiltered references. The resulting classes are representative of the 2 different lattice configurations. Particles in class 2 (box centered on inter-dimer interaction) were 3D refined to 3.09 Å prior to CTF refinement using per-particle defocus, per-micrograph astigmatism, beamtilt estimation, and trefoil estimation. CTF-refined particles were then 3D refined to 2.99 Å. Final halfmaps were exported and sharpened with the resolve cryo-EM^45^ job in phenix^46^ using the “density_modify_unsharpened_maps = True” and “final_scale_with_fsc_ref = False” options, yielding a density map of 2.84 Å. Structure validation is shown in **Extended Data Figure 8A**.

### Cryo-EM 2-protofilament homotypic lattice data processing

2- protofilament homotypic lattice processing was performed as previously described^6^ with minor modifications (**Extended Data Figure 7, middle**). 2-protofilament subtracted particles were backprojected to generate reference maps for subsequent refinements. Initial 3D refinement yielded a 3.03 Å map with a blurred α-tubulin and β-tubulin register. To sort the tubulin register, focus masks corresponding to one of the two protofilaments were used to classify each protofilament individually. Particles masked around the left or right protofilament were subjected to supervised 3D classification without alignment, using 2 molmaps from ChimeraX^44^ as unfiltered 6 Å references - corresponding to the 2 possible tubulin registers - and a regularisation parameter T of 200. This was done for each protofilament using the same starting stack of particles. Particles.star files from left and right protofilament particles in class 2 (box centered on inter-dimer interaction) were then combined using relion_star_combine. The resulting particle stack contained 1,148,862 particles, which refined to 3.03 Å and exhibited a clear homotypic lattice. These particles were then CTF refined using per-particle defocus, per-micrograph astigmatism, beamtilt estimation, and trefoil estimation. CTF-refined particles were 3D refined to 2.93 Å. Final halfmaps were exported and sharpened with the resolve cryo-EM^45^ job in phenix^46^ using the “density_modify_unsharpened_maps = True” and “final_scale_with_fsc_ref = False” options, yielding a density map of 2.79 Å. Structure validation is shown in **Extended Data Figure 8B**.

### Cryo-EM 2-protofilament heterotypic lattice data processing

2-protofilament heterotypic lattice processing was performed as previously described^6^ with minor modifications (**Extended Data Fig. 7, right**). 2-protofilament subtracted particles were backprojected to generate reference maps for subsequent refinements. Initial 3D refinement yielded a 3.03 Å map with a blurred α-tubulin and β-tubulin register. To sort the tubulin register, focus masks corresponding to one of the two protofilaments were used to classify each protofilament individually. Particles masked around the left or right protofilament were subjected to supervised 3D classification without alignment, using 2 molmaps from ChimeraX^44^ as unfiltered 6 Å references - corresponding to the 2 possible tubulin registers - and a regularisation parameter T of 200. This was done for each protofilament using the same starting stack of particles. Particles.star files from left protofilament particles in class 2 (box centered on inter-dimer interaction) and right protofilament particles in class 1 (box centered on heterotimer) were then inner joined using a custom Python script. The resulting particle stack contained 397,803 particles, which refined to 3.35 Å and exhibited a clear heterotypic lattice. These particles were then CTF refined using per-particle defocus, per-micrograph astigmatism, beamtilt estimation, and trefoil estimation. CTF-refined particles were 3D refined to 3.26 Å. Final halfmaps were exported and sharpened with the resolve cryo-EM^45^ job in phenix^46^ using the “density_modify_unsharpened_maps=True” and “final_scale_with_fsc_ref=False” options, yielding a density map of 3.07 Å. Structure validation is shown in **Extended Data Figure 8C**.

### Model building and refinement

Models from a 1-protofilament GDP microtubule structure^6^ were used as starting models. These models were rigid-body fit into each density map using ChimeraX^44^. Models were then iteratively refined into the final density maps using real-space refinement in Phenix (version 1.21.1-5286) and Coot (version 1.31), prioritizing density fit, Ramachandran outliers, geometry, and rotamer outliers in that order. GTP, GDP, and taccalonolide-AJ ligands were added and fit in Coot. Ligand restraints for the covalent linkage between taccalonolide-AJ and βAsp224 was generated using AceDRG^47^ through the CCP4^48^ graphical interface.

### Per-residue structural variance

Per-residue structural variance was calculated as previously described^6^ by taking the variance of pairwise distances for each residue from the difference-distance matrices comparing states. Difference-distance matrices were calculated by measuring the pairwise distances of alpha carbons within a given chain and state and subtracting the corresponding pairwise distances of the same chain in a separate state. The per-residue structural variance for each residue is shown in **Figure. 1C** and **2D**. Our analysis was done in R with help from the bio3D library^49^, although this can be done in many ways. A script that runs this analysis can be found in the Github report associated with this publication.

### Axial twist and protofilament angle calculations

Axial microtubule twists were calculated from the unsharpened, whole-microtubule map in ChimeraX using the following command: >measure symmetry #<volume-spec> minimumCorrelation 0.9 helix 82,0,opt. Inter-protofilament angles for 2-protofilament structures analyzed in **Figure 3E & 3F** and **Extended Data Figure 2B & 2C** were calculated from rotational matrices of fitted β-tubulin subunits using the matchmaker command in ChimeraX.

### Model colorization and vectorization

.defattr, .pb, and .bild files for depicting C-α per-residue structural variance coloration and displacement vectorization were generated using a custom Python script.

### Microtubule gliding assay

COS-7 cells were transfected with 0.5 µg RnKIF5C(1-560)-Avi and 0.5 µg HA-BirA using 97 µL of Opti-MEM (Gibco 31985062) and 3 µL of Mirus TransIT-LT1 transfection reagent (MIR2305). One day later, cells were trypsinized with 0.5 mL of Trypsin-EDTA (0.05%) (Gibco 25300062) and pooled into an equal volume of growth medium (Gibco DMEM 11960044, 10% HyClone FetalClone III SH3010903, 1% Gibco Glutamax 35050061). Cells were pelleted at 1,500 x g for 5 minutes at 4°C. Cell pellets were resuspended in growth medium and re-pelleted. Cell pellets were lysed in 25 µL of lysis buffer (25 mM HEPES pH 7.4, 115 mM KOAc, 5 mM NaOAc, 5 mM MgCl_2_, 0.5 mM EGTA, 1% Triton X-100) supplemented with 1 mM PMSF, 1X protease inhibitor cocktail (Sigma P8340) and 1 mM ATP. Lysates were clarified by centrifugation at 18,000 x g for 10 minutes at 4°C. Supernatants were aliquoted and flash frozen in liquid nitrogen and stored at -80°C.

Microtubules were polymerized the day before use as follows: paclitaxel or tacca AJ microtubules were first polymerized with GTP and without stabilizing agents (0.5 µL of 100mM MgCl_2_, 1µL 28mM GTP, 8 µL of 10 mg/mL porcine brain tubulin (Cytoskeleton T240), and 2 µL of 2.5 mg/mL HiLyte 647 porcine brain tubulin (Cytoskeleton TL670M)) by incubating at 37°C for 30 minutes. Then 100 µL of warm BRB80 with 20 µM paclitaxel or 1 µM tacca AJ was added and incubation at 37°C continued for 3 hours. GMPCPP microtubules were polymerized in the same conditions except with 3 µL of 10 mM GMPCPP replacing the GTP, and 100 µL warm BRB80 with no stabilizing agent was added. All microtubule types were then stored overnight at room temperature in the dark until the following day. Before using, microtubules were pelleted at 21,000 x g and resuspended in 100 µL of BRB80 with 10 µM paclitaxel, 1 µM tacca AJ, or no additives, respectively.

Flow cells were made using a glass slide and 18mm coverslip adhered with strips of melted parafilm. All additions to the flow cell were incubated for 3 minutes. All buffers were supplemented with 10 µM paclitaxel or 1 µM tacca AJ for the corresponding microtubule types. First, 1 mg/mL Biotin-BSA (Sigma A8549) was flowed into the flow cell, followed by wash buffer (500 µg/mL casein/BRB80). Then, 0.5 mg/mL NeutrAvidin (Thermo 31000) was flowed in, followed by wash buffer, lysate mix (17 µl BRB80, 3 µl of RnKIF5C-Avi/HA-BirA lysate, 1 µl of 100 mM ATP, 2 µl of 10 mg/mL casein/BRB80), wash buffer, and finally the microtubule mixture (31 µL BRB80, 1 µL of 100 mM MgCl_2_, 1 µL of 100 mM ATP, 2 µL of oxygen scavenger mix (equal parts 1 M glucose, 100 mM DTT, 10 mg/mL glucose oxidase, and 4 mg/mL catalase), and 10 µL of microtubules (diluted 1:30 from microtubule stocks). Chambers were sealed with melted paraffin wax, and samples were imaged immediately on an inverted Nikon Ti-E/B TIRF microscope with a perfect focus system, a 100 x 1.49 NA oil immersion TIRF objective, three 20 mW diode lasers (488 nm, 561 nm, and 640 nm) and EMCCD camera (iXon^+^ DU879; Andor). Image acquisition was controlled with Nikon Elements software programmed for 100 ms exposures, no delay interval, for 300 loops. All samples were imaged at room temperature. Images were analyzed in FIJI/ImageJ. Arc-chord ratio was determined by drawing a line along the microtubule in the first time frame (arc) and dividing by the straight-line distance between the two end points (chord).

### *In vitro* spermine depolymerization assay

Microtubules were polymerized as follows: paclitaxel or tacca AJ microtubules were first polymerized without stabilizing agents (0.5 µL of 100mM MgCl_2_, 1µL 28mM GTP, 6 µL of 15 mg/mL bovine brain tubulin) by incubating at 37°C for 30 minutes. Then 100 µL of warm BRB80 with 10 µM paclitaxel or 1 µM tacca AJ was added and incubation at 37°C continued for 1 hour. Microtubules were then pelleted by centrifugation at 21,000 x g and resuspended in 200 µL of BRB80 with 10 µM paclitaxel or 1 µM tacca AJ.

Flow cells were made using 18mm coverslips that were HCl acid-washed and silanized with hexamethyldisilazane by vapor deposition for 48 hours adhered to a glass slide with strips of melted parafilm. Chambers were first washed for 3 min with recombinant Tuj1 antibody (gift from Dr. Jeff Moore) diluted 1:50 in BRB80, then washed with BRB80, then microtubules were flowed in and given 3-5 minutes to adhered to the antibody, then washed with 10 µM paclitaxel or 1 µM tacca AJ in BRB80 and a final concentration of 30 mM spermine. Microtubules were then imaged at room temperature by interference reflection microscopy on a C-Trap (Lumicks) using the built-in background subtraction function. Images were taken every second for 30 minutes.

### GMPCPP-stabilized microtubule seeds for Interference Reflection Microscopy

Porcine brain tubulin (Tubulin Protein Porcine Brain, Cytoskeleton, CAT #T240) was resuspended in ice-cold BRB80: 80 mM K-PIPES (Millipore Sigma Cat # P6747-1KG), 1 mM MgCl_2_ (Fisher Scientific Cat #AAJ61014EQE), 1 mM EGTA (Fisher Scientific, Cat #502236999) and mixed with biotinylated porcine brain tubulin (Tubulin protein (biotin) porcine brain, Cytoskeleton, CAT #T333P) on ice at ∼20% biotinylated tubulin and ∼80% unbiotinylated tubulin. Seed stocks were aliquoted and flash-frozen in liquid N_2_, and stored at -80°C until use. On the day of the experiment, seed stocks were diluted to 20 µM tubulin in ice-cold 1X BRB80 (80 mM K-PIPES, 1 mM MgCl_2_, 1 mM EGTA) supplemented with 0.5 mM GMPCPP (Axxora, CAT #NC1303776), and incubated on a heat block at 37°C until use. Immediately before use in a flow chamber, seed stock was sheared by rapid pipetting (20-30 times with a P10 set to 7 μL), and then diluted 1:100 in 1X BRB80 warmed to 37°C and supplemented with GMPCPP.

### Microscopy image acquisition with Interference Reflection Microscopy

Coverslips (22 x 22 mm^2^ Fisher Scientific) were cleaned overnight in 10% 7X Detergent solution (MP Biomedicals, diluted in MilliQ water), rinsed three times with MilliQ water, oxygen-plasma cleaned for 5 min (Harrick Plasma cleaner), and incubated overnight in a vacuum-sealed desiccator with 100 μL of silane (1H,1H,2H,2H-Perfluorodecyltrichlorosilane, Fisher Scientific, CAT #AAL1658406) to generate hydrophobic coverslips. Silanized coverslip hydrophobicity was confirmed via droplet test using MilliQ water and stored in a sealed dehumidification cabinet until use.

Flow chambers were assembled with microscope slides (Fisher Scientific, Fisherbrand Premium Frosted) that were cleaned overnight in 10% MP Biomedicals 7X Detergent (diluted in MilliQ water). Slides were rinsed with MilliQ water, 70% ethanol, then MilliQ water and dried with filtered air and lens paper. Three narrow pieces of double-sided tape (Scotch 665 Permanent Double-sided tape) were affixed to the slide, and then covered with a silanized hydrophobic coverslip, creating two flow chambers per slide. Chambers were prepared at room temperature with all solutions at room temperature: 10 µg/mL NeutrAvidin (Thermo Fisher Scientific, Cat #PI31000) dissolved in 1X BRB80 (80 mM K-PIPES, 1 mM MgCl_2_, 1 mM EGTA) for 10 min, 1% Pluronic 127 (F127, Invitrogen, Cat #P6866) diluted in 1X BRB80 for 10 min, wash with 1X BRB80. Biotinylated microtubule seeds were flowed and incubated at 37°C for up to 10 minutes; seed attachment was monitored on the microscope until 15-20 seeds were attached in the field of view. Chambers with seeds were incubated on a heat block at 37°C until use. Porcine brain tubulin, snap-frozen in liquid nitrogen, was rapid-thawed and placed on ice until use. Ice-cold polymerization solution (80 mM K-PIPES, 1 mM EGTA, 2 mM MgCl_2_, 2 mM GTP (Millipore Sigma Cat #G8877-1G), 0.1 mg/mL BSA (Millipore Sigma Cat #A7030-10G), and 0.1% methylcellulose (Millipore Sigma, Cat #M0512-100G), containing tubulin, was flowed into the chamber, which was subsequently sealed with Biotium CoverGripTM Coverslip Sealant (Fisher Scientific, Cat #NC0154994) and equilibrated at 37°C for 2 minutes.

### IRM image acquisition & image processing

Microtubule movies were captured using Interference Reflection Microscopy (IRM) with a red LED (Nikon) at 20% intensity on a Nikon iLAS2 microscope, using a temperature-controlled stage incubator to maintain a constant temperature at 37°C. Frames were captured with 100 ms exposures at 0.29 sec intervals (3.4 frames per second), for 8-20 minutes, depending on the sample. Focus was maintained using Perfect Focus with shutter control using NIS Elements.

IRM image processing was done with FIJI (NIH^50^). Microtubule movies were flat-fielded to correct for uneven illumination. An out-of-focus image stack (50 frames) was averaged, converted to 32-bits, then normalized by dividing the intensity at each pixel by the average intensity of pixels across the averaged image. Each frame of the microtubule movie was divided by the normalized 32-bit background image. Contrast was further improved by creating a duplicate movie stack, applying a Gaussian blur filter (sigma 20, unscaled units) to the duplicate stack, then dividing the microtubule movie by the blurred stack and creating a 32-bit image. After background collection and contrast improvement, movies were converted to 8-bit images and drift corrected with the Manual Registration plugin^51^, with secondary correction using the Template Matching^52^ plugin if needed. The maximum length reached by each microtubule was visualized using a minimum Z-projection. Using the grouped Z-project function, movies were averaged in groups of 5 frames. Kymographs were generated in batches using the MultiKymograph plugin^53^. Growth and shrinking events were manually traced and analyzed using Microsoft Excel and GraphPad Prism.

### *In vitro* SiR-Tubulin competition assay

Microtubules were polymerized with 0.5 µL of 100mM MgCl_2_, 4 µL of 15 mg/mL bovine brain tubulin, and 2 µL of 2.5 mg/mL HiLyte 488 porcine brain tubulin (Cytoskeleton TL488M) with 1µL 28mM GTP to create GDP-bound microtubule lattice or 3 µL of GMPCPP. Microtubules were incubated for 30 minutes at 37°C, and then 100 µL of warm BRB80 with 1 µM of SiR-Tubulin (Cytoskeleton CY-SC002) was added and microtubules continued to incubate for at least an additional 30 minutes. Before using, microtubules were pelleted at 21,000 x g and resuspended in warm BRB80 with 1 µM SiR-Tubulin and kept at 37°C prior to use.

Flow cells were made using a glass slide and 18mm coverslip adhered with strips of melted parafilm. Chambers were first washed for 3 min with recombinant Tuj1 antibody (gift from Dr. Jeff Moore) diluted 1:50 in BRB80, then blocked with 3 mg/mL BSA in BRB80, then microtubules were flowed in and given 3-5 minutes to adhere, then washed with 1 µM SiR-Tubulin in BRB80. The slide was then brought to the microscope (same as microtubule gliding assays) and an imaging mixture of 1 µM SiR-Tubulin, 5 µM tacca AJ, and oxygen scavengers (equal parts 1 M glucose, 100 mM DTT, 10 mg/mL glucose oxidase, and 4 mg/mL catalase) was added immediately before imaging. Images were taken every 10 seconds for 20 minutes at room temperature. Images were analyzed in FIJI/ImageJ by first applying background subtraction with a rolling ball radius of 20 and then manually drawing 5-pixel-wide lines on microtubules. Intensities were normalized to the average of the first 3 frames.

### In cell fluorescent microscopy

U2OS osteosarcoma cell line modified with a recombination mediated cassette exchange site were maintained in standard culture media (DMEM (Gibco 11960044), 10% Tetracycline-negative screened fetal bovine serum (Gibco A47364-01), 1% Glutamax (Gibco 35050061), and penicillin/streptomycin (Gibco 15140-122))^54^. pEM791 was modified by Gibson assembly to generate donor plasmids with the corresponding fluorescently tagged proteins under the inducible control of a doxycycline promoter. After transfection with donor and Cre recombinase plasmids, cells were given overnight to recover, then cultured in standard media plus 2.5 µg/mL puromycin for 48 hours, then in 5 µg/mL puromycin plus standard media until all cells in negative transfection control were dead and positive colonies were large enough to expand.

Cells were seeded into glass-bottom dishes, and construct expression was induced the day before imaging by adding 1 µg/mL of doxycycline to the media. Immediately before imaging, each dish was washed with 1 mL of Leibovitz’s L-15 media, then 1.5 mL of L-15 media was added to the dish. Vehicle control, paclitaxel, or tacca AJ were diluted into 500 µL of L-15 media for a final concentration of 2.5 µM paclitaxel or tacca AJ once added to the cells. Cells were imaged on an inverted epifluorescence microscope (Nikon Eclipse Ti2) with a 60×, 1.40 NA oil-immersion objective, an ORCA-Flash 4.0LT Plus camera (Hamamatsu), and perfect focus system (Nikon), and maintained and humidified at 37°C with a STX stage-top incubator (Tokai Hit). Cells were equilibrated on the scope for 5-10 min prior to imaging. Images were taken every 10 seconds at a single z-slice for 30 minutes, and media containing vehicle control or treatment was added 30 seconds after imaging began. For overnight imaging, images were taken before treatment, and then after treatment every 10 minutes for 12 hours.

Images were analyzed in FIJI/ImageJ. Microtubule-bound fluorescent intensities were determined by correcting for drift using the built-in plugin “Linear Stack Alignment with SIFT”, manually segmenting cells, and then processing each timepoint by running the built in “Tubeness” plugin with a sigma of 0.1083, converting images to 8-bit and applying a threshold cutoff of 30 to 255, converting to a mask using the built in “Skeletonize (2D/3D)” plugin, and then removing any particles with an area of less than 0.06 µm^2^. EB3 foci were quantified by preparing images of individual cells in the same manner, pre-processing the images by using the “Subtract Background” function with a rolling ball radius of 2, applying the “Remove Outliers” function with a radius of 1 and threshold of 25, and then applying the “Despeckle” function. A threshold was then applied using a cutoff range of 45 to 255 for intensities, the intensities converted to a mask, the “Close-“ function applied to remove small holes in the mask, and then number of particles counted with an area between 0.35 and 0.23 µm^2^ were considered EB3 foci. Values were normalized to the average of the 3 time frames for that cell prior to treatment.

Microtubule stability in cells was assayed by cold-shock by transfecting U2OS acceptor cells with GFP-TUBA1A 48 hours prior to imaging. Each dish was washed with 1 mL of Leibovitz’s L-15 media, then 2 mL of L-15 media was added to the dish containing vehicle control, 2.5 µM of paclitaxel, or 2.5 µM of tacca AJ. Cells were then incubated for 30 minutes after treatment, then imaged before the cold shock. All three conditions were simultaneously placed at -20°C for 10 minutes, then imaged at 37°C within 15 minutes.

## Data Availability

Coordinates for 1-protofilament, 2-protofilament homotypic, and 2-protofilament heterotypic tacca AJ-GDP microtubule structures are available at the Protein Data Bank (PDB) with accession codes XXXX, XXXX, and XXXX, respectively. Cryo-EM density maps for 1-protofilament, 2-protofilament homotypic, and 2-protofilament heterotypic tacca AJ-GDP microtubule structures are available at the Electron Microscopy Data Bank (EMDB) with accession codes XXXX, XXXX, and XXXX, respectively. The cryo-EM dataset for tacca AJ-GDP microtubules is available at the Electron Microscopy Public Image Archive (EMPIAR) with accession code XXXX.

## Code Availability

Detailed cryo-EM processing pipeline and scripts for cryo-EM processing and structural analysis are available at GitHub (https://github.com/cianfrocco-lab/nvangos).

**Extended Data Figure 1 - Comparison of longitudinal conformational changes between tacca AJ, GDP, and paclitaxel.** (A) Homotypic and (B) heterotypic lattice comparisons shown with coloring based on longitudinal differences between indicated structures.

**Extended Data Figure 2 - Comparison of lateral changes between tacca AJ, GDP, and paclitaxel.** (A) Schematic of microtubule structure. (B) Homotypic and (C) heterotypic lattice comparisons.

**Extended Data Figure 3 - Microtubule gliding, spermine treatment, and SiR-tubulin assays. (**A) Microtubule lengths analyzed for tortuosity index in Figure 4B. (B) Gliding velocity of microtubules analyzed in Figure 4A-C. Outlined points represent the median from each technical replicate. Error bars are the mean ± 95% confidence interval of technical replicates. (C) Example kymographs of unlabeled paclitaxel or tacca AJ microtubules observed by interference reflection microscopy treated with 30 mM spermine. Cartoons below each kymograph help depict the ends of the microtubules. Scale bar height = 10 min, width = 2 µm. (D) Example kymographs of SiR-tubulin on GMPCPP or GDP microtubules after washing the chamber with vehicle control buffer. Scale bar height = 2 min, width = 1 µm. (E) Quantification of normalized SiR-tubulin intensity on GMPCPP microtubules after washing the chamber with vehicle control buffer. Mean ± 95% confidence interval, n = 48 microtubules.

**Extended Data Figure 4 - Tacca AJ quickly disrupts MT dynamics in cells.** (A) Example time point images of EB3-mStayGold intensity (inverted greyscale) in live U2OS cells after treatment. Dotted red boxes depict 5X zoom. (B) Quantification of EB3-mStayGold foci after treatment, normalized to the number of foci before treatment. Line = mean, shaded area = 95% confidence interval. Control n = 19 cells; paclitaxel = 15 cells; tacca AJ = 16 cells.

**Extended Data 5 - Tacca AJ-stabilized MTs remain compacted for 12 hours.** Example time point images of MAP2-mStayGold or DCX-mStayGold intensity (inverted greyscale) in live U2OS cells after treatment.

**Extended Data Fig. 6 - Tacca AJ whole-microtubule cryo-EM data processing workflow.** Colored numbers in parentheses indicate particle number, class number, or resolution for a given experimental condition.

**Extended Data Fig. 7 - Tacca AJ 13-protofilament microtubule cryo-EM data processing workflow.** (continues from Extended Data Fig. 6) Cryo-EM processing workflows for 13-protofilament tacca AJ microtubules, split into 1-protofilament (left, light orange box), 2-protofilament homotypic (middle, orange box), and 2-protofilament heterotypic (right, dark orange box) lattice structures.

**Extended Data Fig. 8 - Tacca AJ 13-protofilament structure validation.** FSC curves for unsharpened (top) and density-modified (bottom) maps, final density map colored by local resolution, fitted model colored by b-factor, 3D particle orientation distribution, and zoom-in density comparisons for 1-protofilament (A), 2-protofilament homotypic (B), and 2-protofilament heterotypic (C) lattice structures.

