## Supplemental Figures for "Taccalonolide AJ uncouples tubulin GTP hydrolysis from microtubule depolymerization"

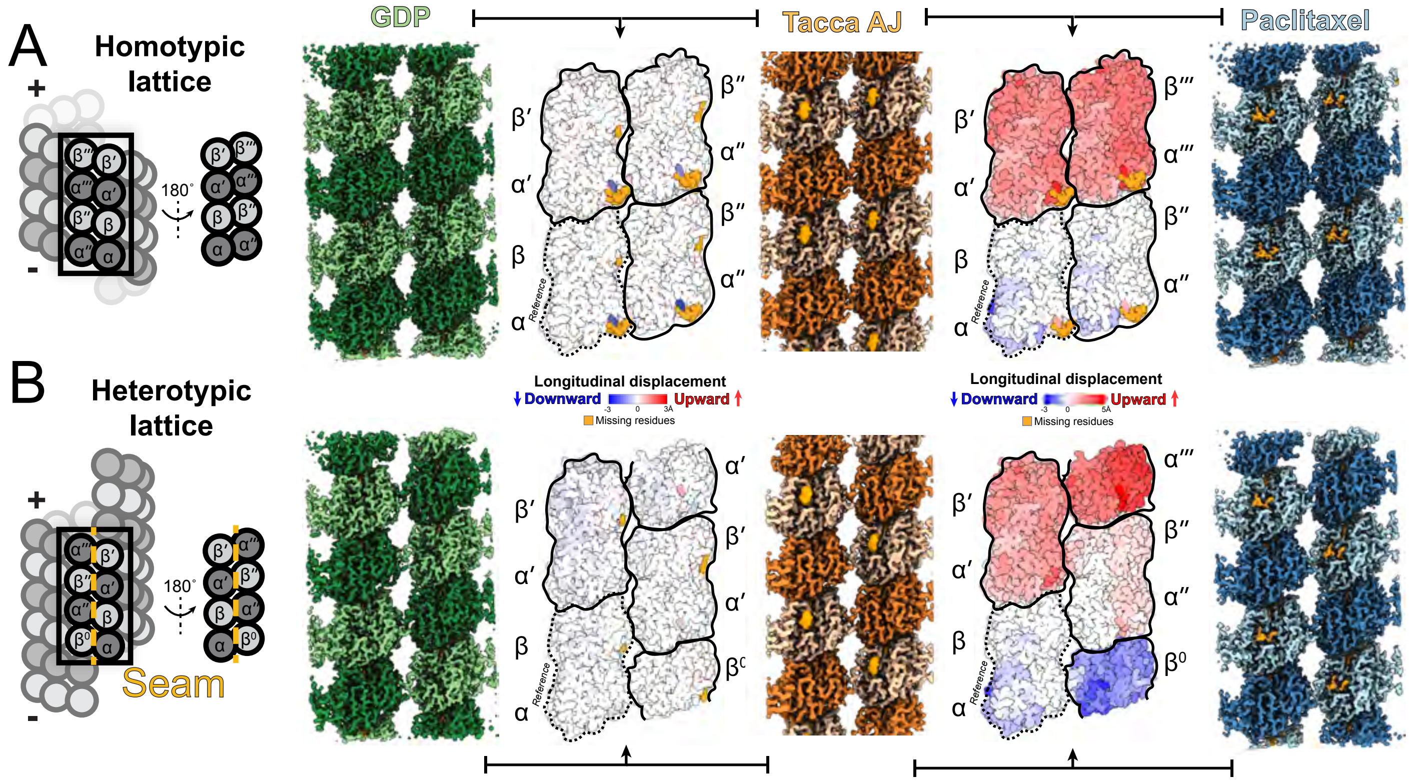


Extended Data Figure 1 - Comparison of longitudinal conformational changes between tacca AJ, GDP, and paclitaxel.

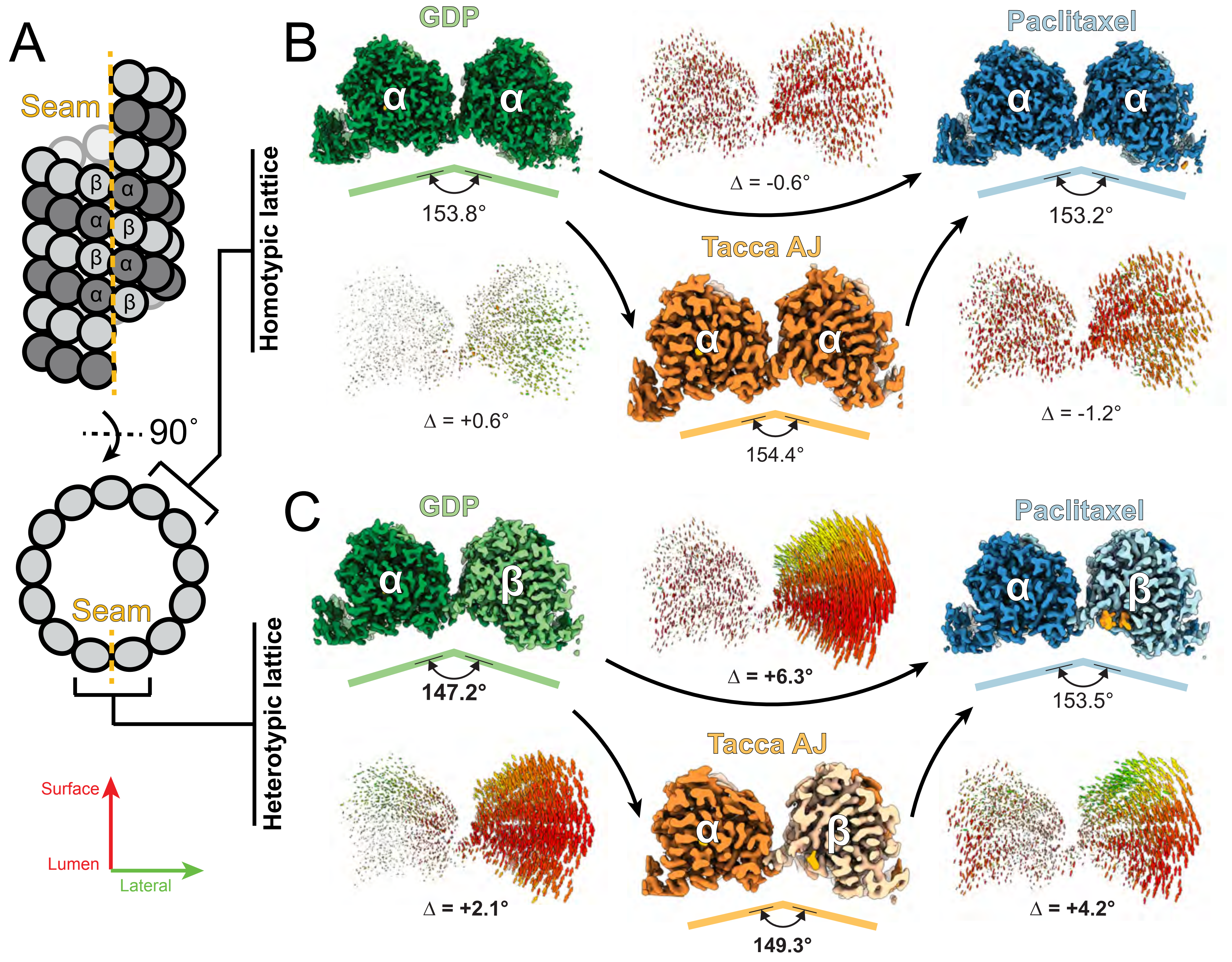


Extended Data Figure 2 - Comparison of lateral changes between tacca AJ, GDP, and paclitaxel.

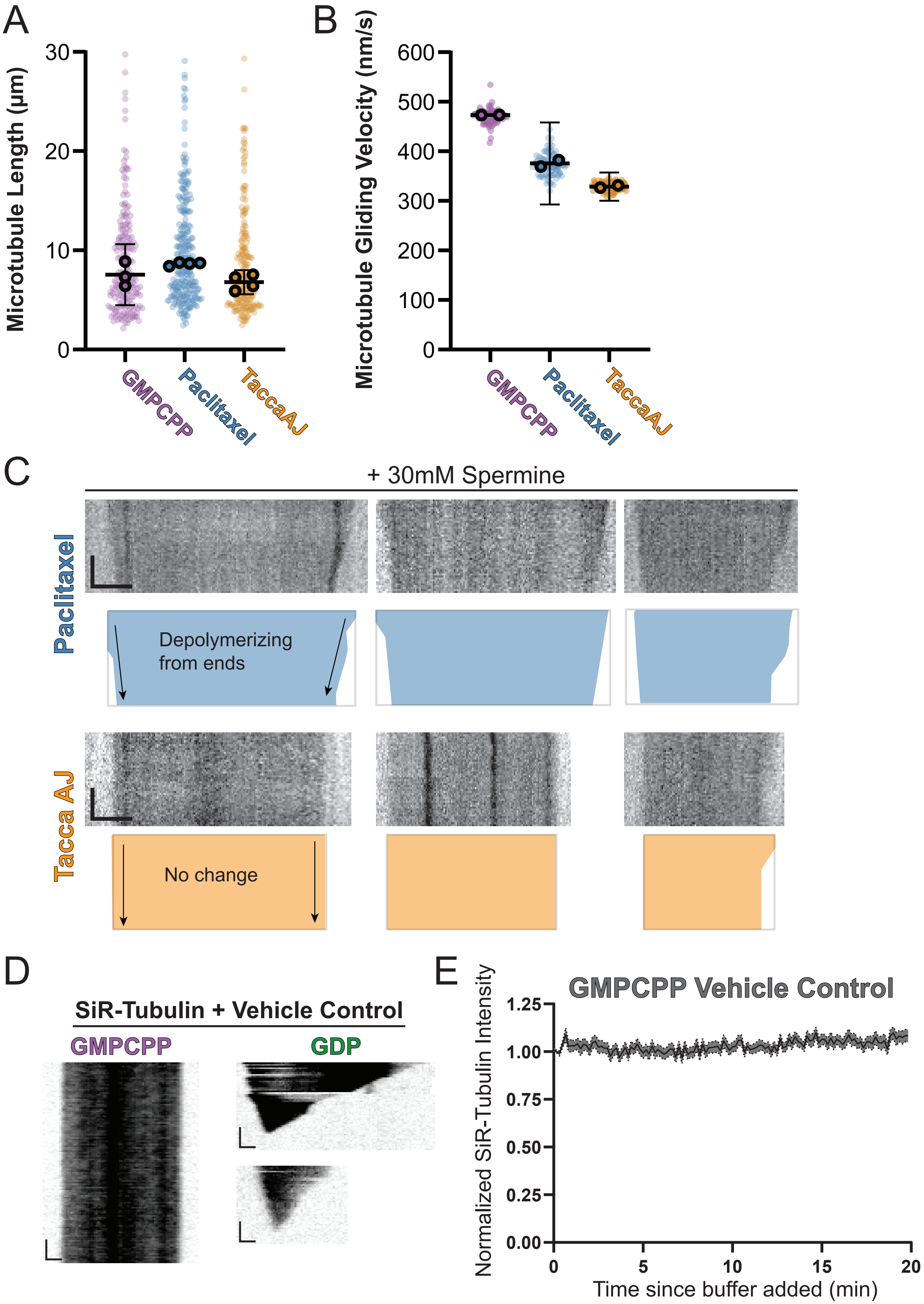


Extended Data Figure 3 - Gliding assay, spermine treatment, and SiR-Tubulin control

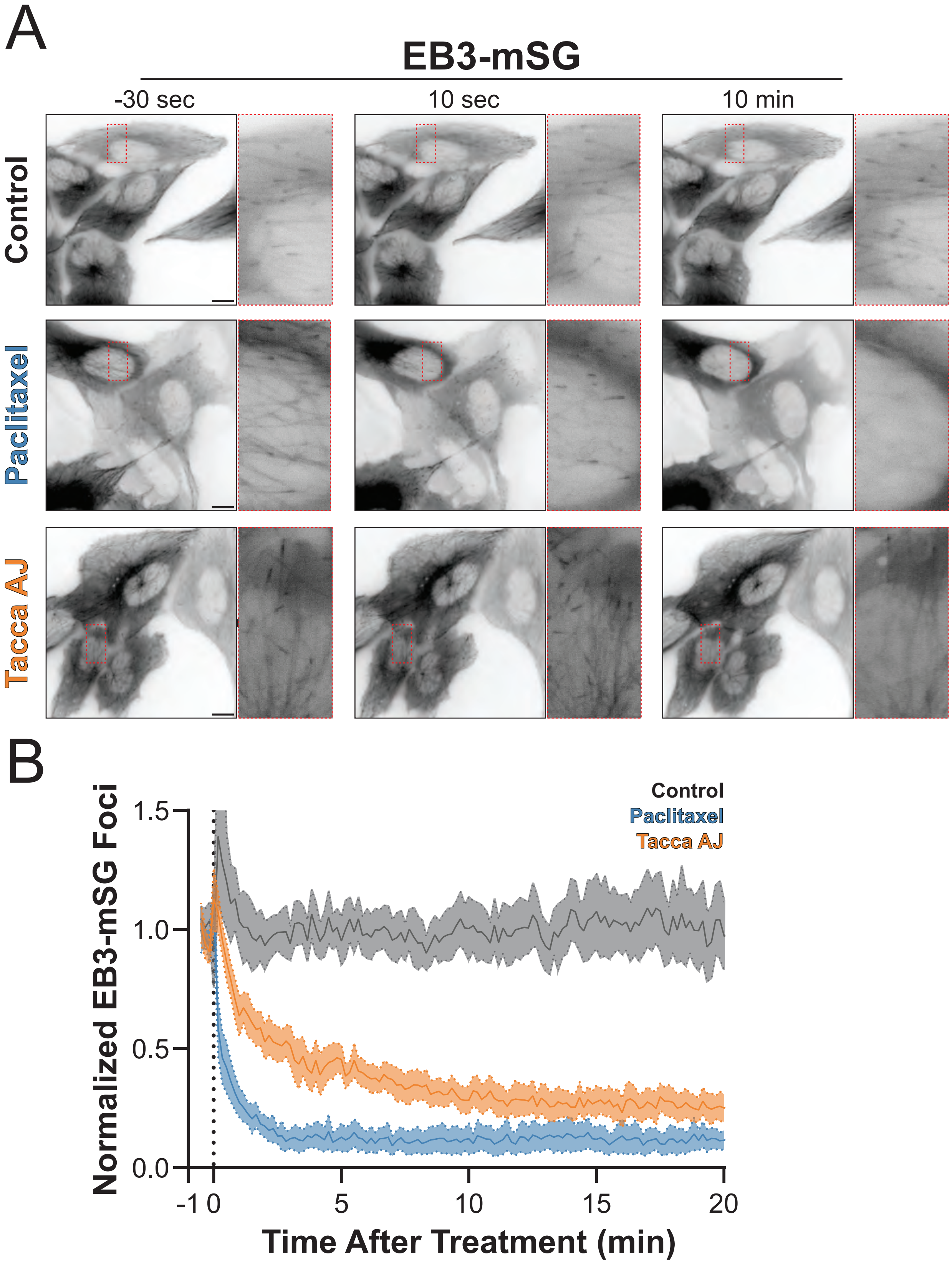


Extended Data Figure 4 - Tacca AJ quickly disrupts MT dynamics in cells

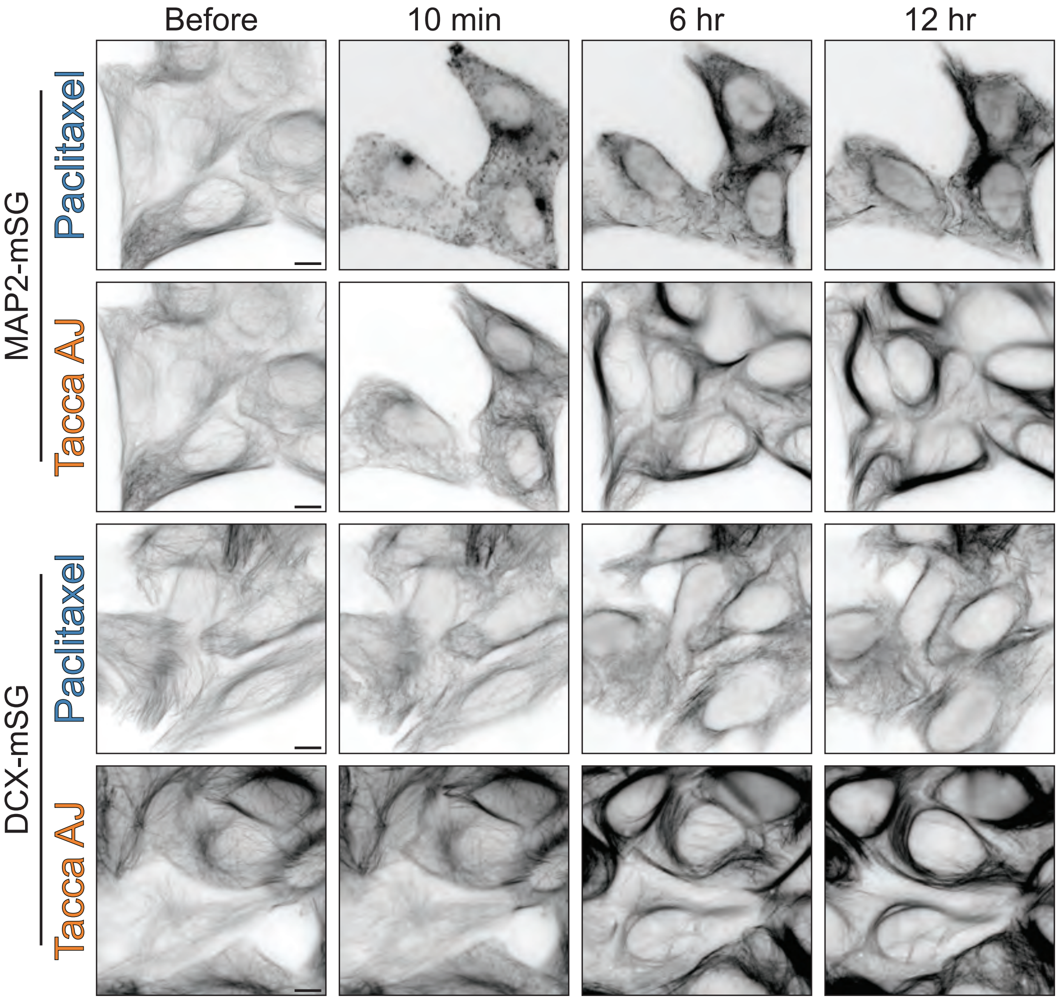


Extended Data Figure 5 -Tacca AJ-stabilized MTs remain compacted for 12 hours

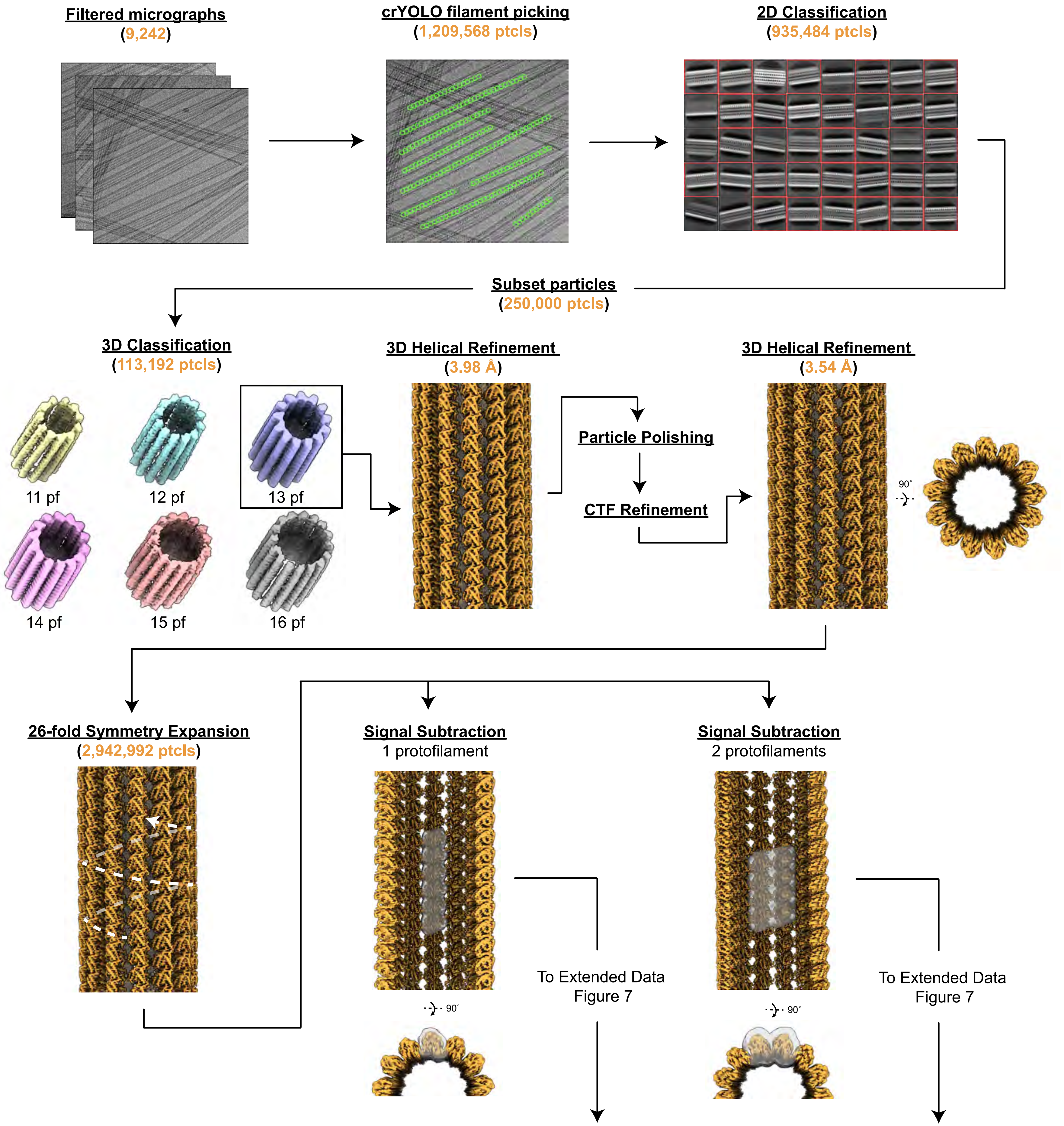


Extended Data Figure 6 - Tacca AJ whole-microtubule cryo-EM data processing workflow.

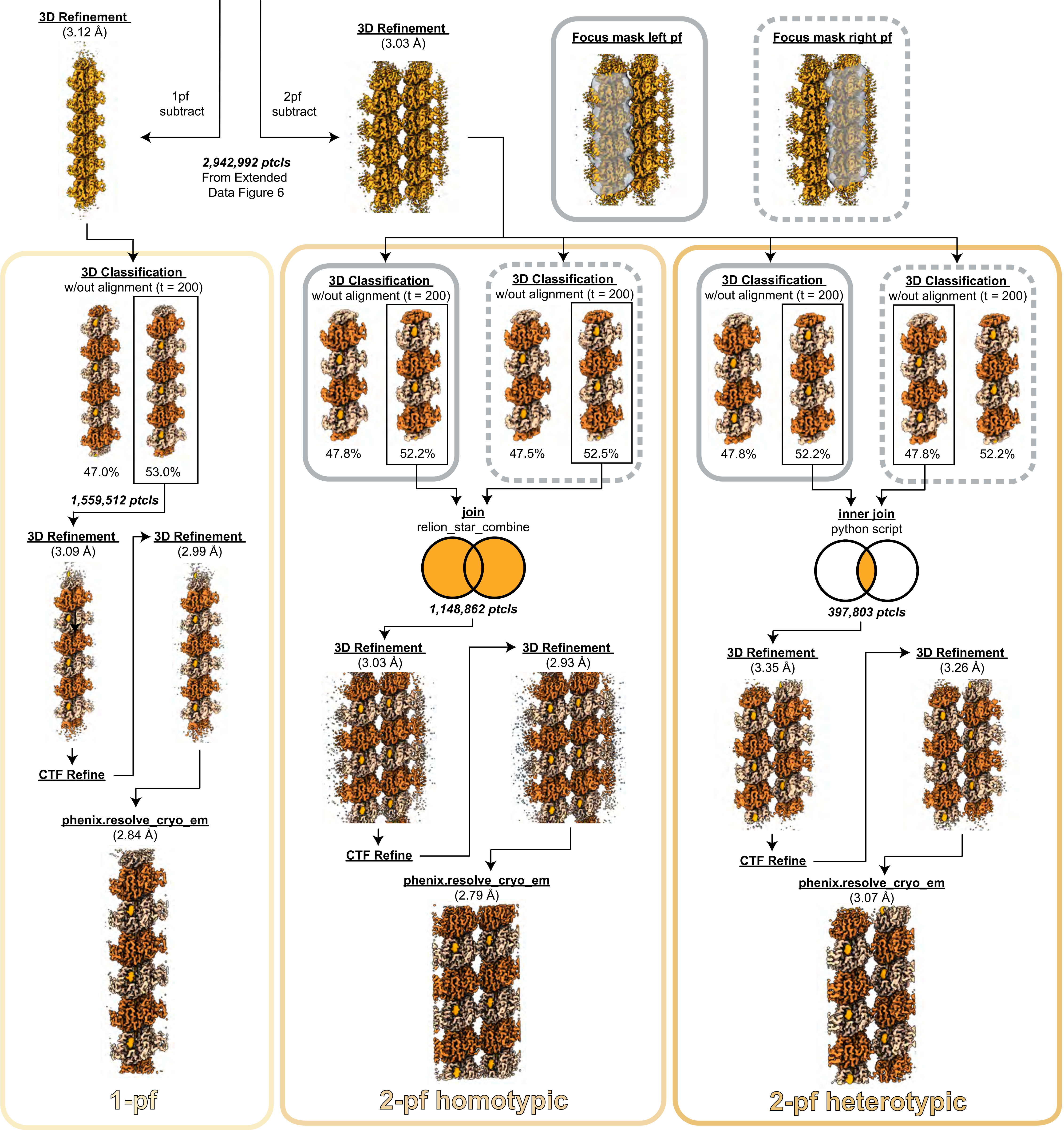


Extended Data Figure 7 - Tacca AJ 13-protofilament microtubule cryo-EM data processing workflow.

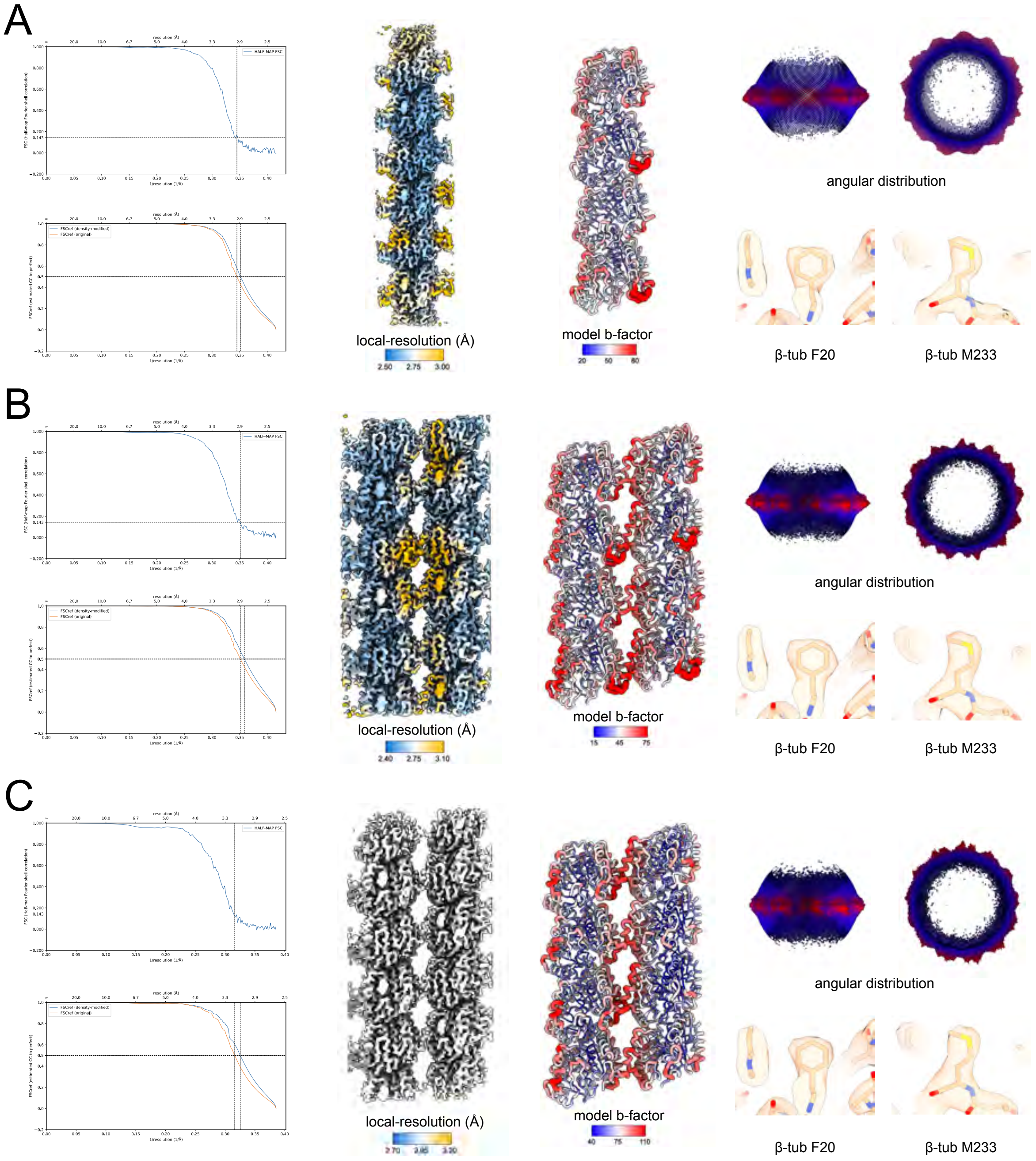


Extended Data Figure 8 - Tacca AJ 13-protofilament structure validation.

| | Tacca-GDP MT | | |
|---|---|---|---|
| **Data collection and processing** | 1-pf | 2-pf (homotypic) | 2-pf (heterotypic) |
| **PDB ID** | XXXX | XXXX | XXXX |
| **EMDB ID** | XXXXX | | |
| Microscope | Glacios | | |
| Voltage (kV) | 200 | | |
| Detector | Falcon 4 | | |
| Electron exposure ($e^-/Å^2$) | 52 | | |
| Frame number | 55 | | |
| Calibrated pixel size (Å) | 0.88838 | | |
| Defocus range (µm) | 0.5-2.0 | | |
| Symmetry imposed | C1 | | |
| Movies | 9,602 | | |
| Initial particle images (before symmetry-expansion) | 1,209,568 | | |
| Final particle images | 1,558,512 | 1,148,862 | 397,803 |
| Map resolution (Å) | 2.99 | 2.93 | 3.26 |
| FSC threshold | 0.143 | 0.143 | 0.143 |
| Map sharpening factor ($Å^2$) | -90.2 | -86.1 | -90.4 |
| Density-modified map resolution (Å) | 2.84 | 2.79 | 3.07 |
| Density-modified FSC threshold | 0.5 | 0.5 | 0.5 |
| Helical symmetry (13-pf MT map) | | | |
| Single subunit rise (Å) | 9.31 | | |
| Helical twist (°) | -27.7 | | |
| Dimer rise (Å) | 80.54 | | |
| Axial twist (°) | -0.11 | | |
| **Refinement** | | | |
| Initial models (AlphaFold ID) | | | |
| α-tubulin | PDB 35SN | | |
| β-tubulin | PDB 35SN | | |
| Model resolution (Å) | 2.51 | 2.47 | 2.68 |
| FSC threshold | 0.5 | 0.5 | 0.5 |
| Model composition | | | |
| Non-hydrogen atoms | 13,774 | 27,548 | 27,280 |
| Protein residues | 1,730 | 3,460 | 3,420 |
| Ligands | 8 | 16 | 12 |
| B factors ($Å^2$) | | | |
| Proteins | 47.33 | 45.28 | 69.76 |
| Ligands | 49.68 | 43.91 | 69.52 |
| R.M.S. deviations | | | |
| Bond lengths (Å) (# > 4 ) | 0.003 (0) | 0.003 (0) | 0.004 (0) |
| Bond angles (°) (# > 4 ) | 0.520 (2) | 0.519 (4) | 0.569 (7) |
| Validation | | | |
| MolProbity score | 1.81 | 1.80 | 2.09 |
| Clashscore | 7.69 | 7.91 | 9.59 |
| Poor rotamers (%) | 2.65 | 2.41 | 4.19 |
| Ramachandran plot | | | |
| Favored (%) | 97.68 | 97.65 | 97.41 |
| Allowed (%) | 2.32 | 2.35 | 2.59 |
| Disallowed (%) | 0.00 | 0.00 | 0.00 |
| Rama-Z score | 0.76 | 1.01 | 0.88 |

Table 1 - Cryo-EM data collection, processing, and refinement statistics.
